# TLR2-mediated microbial sensing by intestinal stem cells coordinates epithelial antimicrobial defense

**DOI:** 10.64898/2026.05.04.722581

**Authors:** Aviya Habshush Menachem, Sacha Lebon, Christoph Killian, Vladyslav Holiar, Avital Sarusi Portuguez, Natalia Davidzohn, Barak Toval, Carmel Sochen, Smadar Levin-Zaidman, Nili Dezorella, Leilah Otikovs, Deborah Duran, Ilan Kent, Myriam Grunewald, Lorenz Adlung, Moshe Biton

**Affiliations:** Department of Immunology and Regenerative Biology, Weizmann Institute of Science, Rehovot, Israel; First Department of Medicine, University Medical Center Hamburg-Eppendorf, Hamburg, Germany; The Mantoux Bioinformatics institute of the Nancy and Stephen Grand Israel National Center for Personalized Medicine, Weizmann Institute of Science.; Department of Chemical Research Support, Weizmann Institute of Science, Rehovot, Israel; The Hadassah Organoid Center, the Hadassah-Hebrew University Medical Organization, Jerusalem, Israel; Faculty of Medical and Health Sciences, School of Medicine, Tel-Aviv University, Tel Aviv 69978, Israel; Department of General Surgery, Chaim Sheba Medical Center, Ramat Gan 52621, Israel; Hamburg Center for Translational Immunology (HCTI) and Center for Biomedical AI (bAIome), University Medical Center Hamburg-Eppendorf, Hamburg, Germany

## Abstract

Intestinal regeneration and host defense require adaptation to environmental cues, but the mechanisms underlying this coordination remain unclear. We show that intestinal Lgr5⁺ stem cells act as luminal sensors via apically localized Toll-like receptor 2 (TLR2), enabling direct detection of microbiota-derived signals. We identify apical TLR2 activation as a mechanism of luminal sensing in adult stem cells and show that it controls epithelial differentiation, antimicrobial peptide production, and crypt organization, with a particularly strong influence on Paneth cell maturation. Genetic ablation of constitutive, epithelial, or stem cell-specific TLR2 disrupts these processes, leading to impaired antimicrobial defense and altered epithelial composition. Using germ-free mice and human intestinal organoids, we demonstrate that this pathway is microbiota-dependent and evolutionarily conserved, respectively. These findings support a model in which stem cells act as active integrators of environmental information and suggest a broader principle by which barrier tissues couple microbial sensing to regeneration and host protection.

## Introduction

The mucosal surface of the gastrointestinal (GI) tract is composed of a single layer of intestinal epithelial cells (IECs) that forms a critical interface between the host and the external environment. Under physiological conditions, the intestinal epithelium integrates signals derived from nutrients and commensal organisms to maintain a balanced immune response^1,2^. Disruption of this balance is associated with intestinal pathologies, including inflammatory bowel disease (IBD)^1,3^. Accordingly, IECs are increasingly recognized not only as physical barriers but also as active participants in sensing and responding to environmental stimuli^4–9^.

All epithelial lineages of the intestine are derived from Lgr5⁺ intestinal stem cells (ISCs), which reside at the base of crypts and continuously replenish the epithelium^10,11^. These stem cells give rise to specialized cell types with distinct roles in host defense, including mucus-secreting goblet cells, enterocytes that produce specific antimicrobial peptides (AMPs), such as the Reg3 family^4,9,10^, and Paneth cells, which both support the stem cell niche and secrete a broad repertoire of AMPs^1,12,13^. Together, these differentiated lineages maintain epithelial integrity and regulate host-microbiota interactions. However, the mechanisms by which antimicrobial programs are established during epithelial replenishment remain poorly understood.

ISCs have traditionally been viewed as passive immunological cells, with protection of the stem cell compartment provided by their surrounding niche cells, including Paneth cells that secrete AMPs to limit microbial access to the stem cell compartment^12,14^. Nevertheless, accumulating evidence suggests that ISCs can actively respond to environmental cues^15–18^. Under stress or infection, ISC-driven epithelial remodeling rapidly alters epithelial lineage allocation to promote host defense^4,15^. In particular, recent work from our group has shown that ISC-intrinsic inflammasome activation during *Salmonella* intracellular infection drives differentiation toward AMP-producing lineages^19^. These observations raise the possibility that ISCs may also directly sense luminal microbial signals and regulate epithelial regeneration in response to environmental inputs.

Central mediators of host-microbe interactions are the pattern recognition receptors (PRRs), including Toll-like receptors (TLRs) and NOD-like receptors (NLRs)^20–22^. These PRRs are critical for maintaining intestinal homeostasis and are implicated in gut pathologies such as IBD and colorectal cancer (CRC) ^23–27^. Importantly, TLR2 is a cell-surface PRR that recognizes conserved and diverse microbial, viral, and fungal ligands and plays a pivotal role in mucosal immunity^27,28^. While TLR signaling in differentiated epithelial cells has been extensively characterized, their role within ISCs *in vivo* remains poorly defined. In particular, whether ISCs directly detect microbial signals and how such sensing influences stem cell fate and epithelial differentiation remain key unresolved questions.

Here, we investigate how ISCs integrate microbial cues to regulate epithelial composition and antimicrobial defense. We identify TLR2 as an ISC-enriched PRR and show that Lgr5⁺ ISCs can sense luminal microbiota via apical TLR2 activation. This sensing mechanism enables ISCs to couple microbial signals to epithelial differentiation, promoting Paneth cell maturation and the establishment of antimicrobial programs. Using *in vivo* genetic models, stem-enriched *ex vivo* spheroid, monolayer cultures, and human organoids, we demonstrate that Lgr5+-ISC-specific TLR2 activation initiates an epithelial AMP program and shapes epithelial remodeling in homeostasis, orchestrating the delicate balance of host-microbiota interactions in the gut. Together, our findings suggest that adult stem cells can function as microbial sensors and uncover a mechanism by which environmental signals are integrated to coordinate regeneration and host defense.

## Results

### TLR2 is expressed in ISCs under homeostasis

A hallmark of intestinal stem cells (ISCs) is their ability to rapidly remodel the epithelium in response to stress^4,9,15,29^. Because epithelial turnover occurs within 5 days or less, Lgr5^+^-ISC-driven remodeling enables rapid adaptation to luminal cues. However, how Lgr5^+^-ISCs sense luminal content remains unclear. To address this, we searched for pattern recognition receptors (PRRs) enriched in Lgr5^+^-ISCs using our previously generated high-definition full-length single cell RNA-sequencing (scRNA-seq) dataset^29^ of Lgr5^+^-ISCs and all other major epithelial subsets from the Lgr5-EGFP-IRES-Cre^ERT2^ (Lgr5-GFP) mouse model^30^, focusing on Toll-like receptors (TLRs), which sense extracellular microbe-associated molecular patterns (MAMPs). Among murine TLRs, *Tlr3* mRNA was broadly expressed across epithelial subsets, with the highest levels in tuft cells. In contrast, *Tlr2* was highly enriched in Lgr5^+^-ISCs, decreased progressively in transient amplifying (TA) cells and progenitors, and was nearly absent from differentiated epithelial populations (**Fig. 1a**). To validate *Tlr2* expression under homeostatic conditions, we isolated EpCAM^+^ epithelial cells or Lgr5^+^-ISCs from Lgr5-GFP reporter mice (**Fig. 1b and Extended Data Fig. 1a**). Quantitative PCR (qPCR) confirmed significant enrichment of *Tlr2* expression in Lgr5^+^-ISCs compared with the epithelial compartment (**Fig. 1b**). Single-molecule fluorescence *in situ* hybridization (smFISH) further supported these findings, showing co-localization of *Tlr2* and *Lgr5* RNA molecules in crypt regions of wild-type (WT) mice, but not in TLR2 constitutive knockout (TLR2 cKO) mice (**Fig. 1c**). The small intestine (SI) is anatomically divided into the duodenum, jejunum, and ileum, each with distinct physiological roles. While the duodenum and jejunum are primarily specialized for digestion and nutrient absorption, the ileum serves as a reservoir for commensal bacteria due to its proximity to the large intestine^31^. Consistent with this proximal-to-distal microbial gradient, *Tlr2* expression followed a segmental pattern along SI crypts, with the highest levels in the ileum (**Fig. 1d**). Together, these results indicate that *Tlr2* expression is preferentially enriched in ileal ISCs, consistent with localization to a region of increased microbial exposure.

**Fig. 1.**
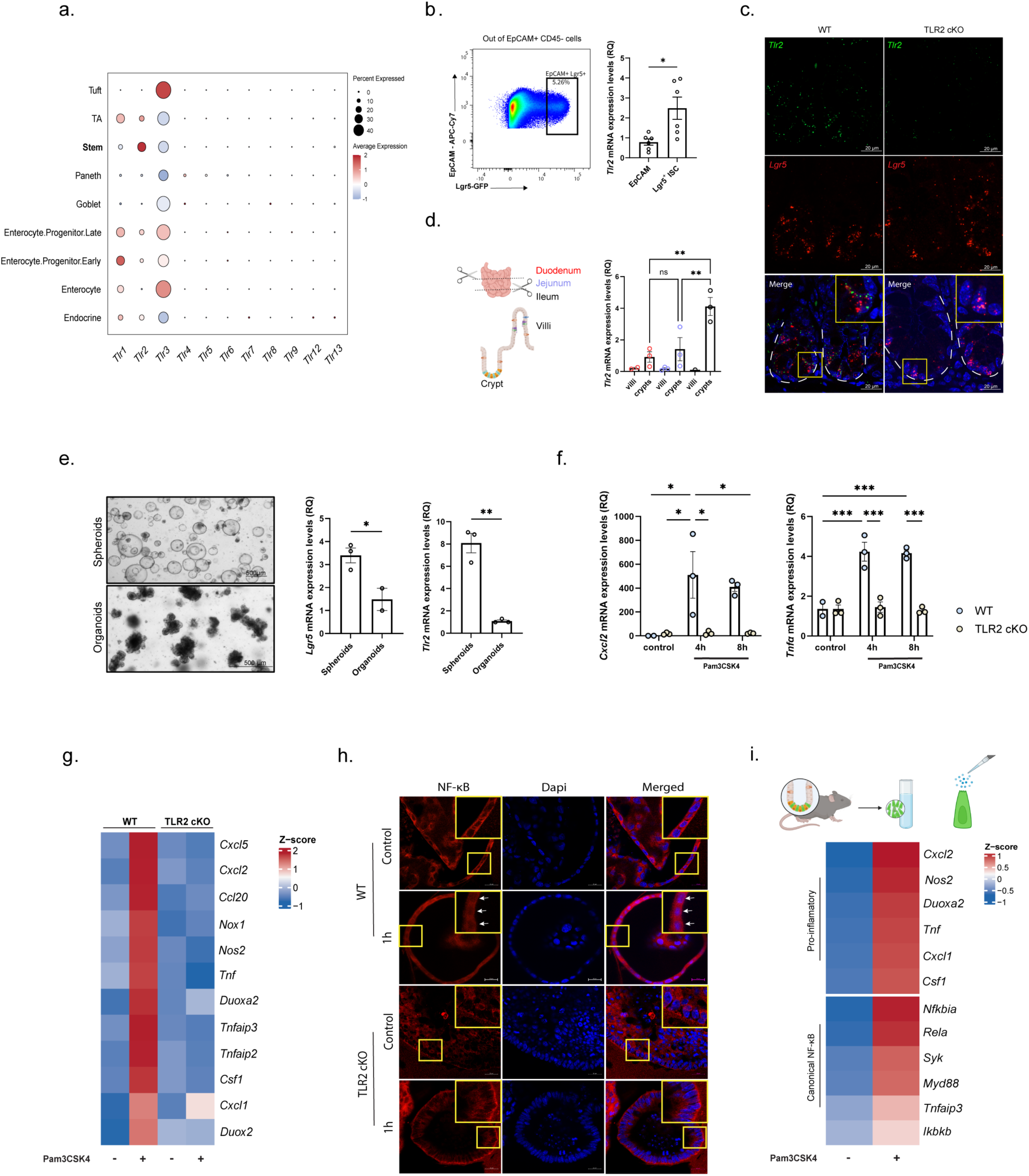
Ileal Lgr5^+^-ISCs express functional TLR2. (**a**) Plate-based epithelial single cell re-analysis of Toll-like receptors (TLRs). Dot plot of 11 murine TLRs expressed in different IECs under homeostasis, taken from a publicly available dataset^1^. The dot size indicates the proportion of cells expressing a gene, and the color indicates the mean expression levels. (**b**) Representative flow-cytometry plot of Lgr5+-ISC population in Lgr5-GFP mouse model (left) and Relative quantification (RQ) of *Tlr2* expression in murine Lgr5^+^ cells under homeostasis using quantitative PCR (qPCR) (Right). Data are presented as mean ± SEM. *n*=6 mice per condition, \**p* <0.05, two-tailed *Student’s t-test*. (**c-d**) *Tlr2* expression on ileal Lgr5^+^-ISC under homeostasis. (c) Co-localization of *Lgr5* and *Tlr2* molecules in ISCs. Single-molecule fluorescence in situ hybridization (smFISH) of *Tlr2* (green) and *Lgr5* (red), co-stained with E-Cadherin and DAPI (blue). E-Cadherin stains IEC boundaries, and DAPI stains nuclei. Scale bar, 20μm. Single channel of each color (red or green) or merged images (lower panels); WT crypt (left panels) and TLR2 KO crypt (right panels). Insets x2 magnification. (d) Experimental scheme of the different segments of the small intestine for villi and crypts isolation (left) and RQ of *Tlr2* expression levels using qPCR (right). Data are presented as mean ± SEM. *n*=3 mice per region; ns (non-significant), \**p* <0.05, \*\**p* <0.01, *one-way ANOVA*. (**e**) Primary mouse SI spheroids and organoids from ileal crypts. Representative pictures of stem-enriched spheroids (upper panel) and differentiated organoids (lower panel) (left). Scale bar, 500μm. RQ of *Tlr2* and *Lgr5* expression levels in spheroids and organoids using qPCR (right). Data are presented as mean ± SEM. *n*=3 biological repetitions for spheroids or 2 for organoids, \**p* <0.05, two-tailed *Student’s t-test*. (**f**) RQ of *Cxcl2* and *Tnfα* expression levels in spheroids generated from WT vs. TLR2 cKO mice after 4h or 8h of *ex vivo* activation with TLR2/TLR1 agonist (Pam3Csk4, 1μg/mL) or without (control) using qPCR. Data are presented as mean ± SEM. *n*=3 biological repetitions, \**p* <0.05, \*\*\**p* <0.001, two-way ANOVA. (**g**) Heatmap of top TLR2 target genes of bulk RNA-seq from WT vs TLR2 cKO stem-enriched spheroids after 4h of activation with or without Pam3Csk4 (1μg/mL). Mean Z score of log_2_(TPM+1), color bar. *n*=3 biological repetitions for WT and 2 for TLR2 cKO. (**h**) NFĸB pathway activation. Representative images of immunofluorescence assay (IFA) of NFĸB subunit (p65, red) and DAPI of stem-enriched spheroids in WT (top panels) or TLR2 cKO (lower panels) at time 0 or 1 hour after Pam3Csk4 induction (1μg/mL). Scale bar, 20μm. Insets, x2 magnification. Arrows show p65 localization to the nucleus (**i**). *Tlr2* activation of Lgr5+-ISC *ex vivo* scheme (upper panel). Heatmap of TLR2 target genes (inflammatory) and canonical NFĸB targets from sorted WT Lgr5+-ISCs after 4h of *ex vivo* activation with or without Pam3Csk4 (1μg/mL) by bulk RNA-seq. Average relative expression (mean Z score of log_2_(TPM+1), color bar). *n*=3 biological repetitions for control and 2 for Pam3Csk4 activation.

### TLR2 is functionally active in intestinal stem cells

The gut contains not only epithelial, stromal, and immune cells, but also diverse microbial communities, including bacteria, fungi, viruses, and parasites^27^, many of which can be detected by cell-surface innate immune receptors such as TLR2^28^. To determine whether TLR2 is functionally active in ISCs, we used long-term *ex vivo* cultures of adult ISCs and differentiated epithelial cells in 3D, namely spheroids and organoids, respectively, derived from WT mice^32,33^ (**Methods**, **Fig. 1e**). As expected, stem-enriched spheroids expressed higher levels of both *Lgr5* and *Tlr2* than differentiated organoids (**Fig. 1e**). We next stimulated spheroids and organoids with the TLR2/TLR1 agonist, Pam3CysSerLys4 (Pam3CSK4), for short time intervals (0, 4, and 8 hours; **Methods**, **Extended Data Fig. 1b**). Pam3CSK4 induced a robust innate immune response in WT spheroids, including upregulation of *Tlr2* itself and of canonical TLR2 target genes such as *Cxcl2* and *Tnfα* after 4 hours, whereas organoids showed only a weak response (**Fig. 1f** and **Extended Data Fig. 1b-c**). This response was absent in TLR2 cKO spheroids, confirming pathway specificity (**Fig. 1f** and **Extended Data Fig. 1c**). TLR2 signaling activates downstream cascades leading to NF-κB activation. Consistent with this, bulk RNA-seq of WT but not TLR2 cKO spheroids revealed induction of inflammatory and NF-κB-associated target genes following Pam3CSK4 stimulation (**Fig. 1g**, **Supplementary Table 1**). At the protein level, Pam3CSK4 treatment induced nuclear translocation of the NF-κB subunit p65 in WT spheroids, but not in TLR2 cKO spheroids (**Fig. 1h** and **Extended Data Fig. 1d**). Importantly, sorted Lgr5^+^-ISCs, when stimulated *ex vivo* with Pam3CSK4 for 4 hours, similarly upregulated TLR2 target genes and NF-κB pathway components (**Fig. 1i**, **Extended Data Fig. 1e**, and **Supplementary Table 2**). These data demonstrate that TLR2 is functionally active in Lgr5^+^-ISCs and can trigger a pro-inflammatory NF-κB transcriptional program.

### Polarized activation of TLR2 on stem-enriched epithelial monolayers

Epithelial cells exhibit apical-basal polarity, which is essential for barrier function and directional signal sensing^26,34,35^. Previous studies reported TLR2 localization at both the apical and basolateral surfaces in murine follicle-associated epithelium, with a lower apical signal in villus epithelium ^36,37^. We therefore asked whether TLR2 signaling in Lgr5^+^-ISCs is triggered from the apical side, enabling detection of luminal MAMPs, or from the basolateral side, to detect pathogenic bacteria, as observed in TLR9-expressing colorectal cell lines, HCA-7 and Caco-2 ^26^. To address this, we established long-term two-dimensional (2D) stem-enriched monolayers from SI spheroids using conditions adapted from colon cultures^38,39^ (**Methods** and **Fig. 2a**). These cultures formed a polarized single-cell epithelial layer, confirmed by immunostaining for Villin1 and CD138, which mark the apical and basolateral membranes, respectively^39^ (**Fig. 2b**). Monolayer integrity was validated by transepithelial electrical resistance (TEER) measurements and FITC-dextran permeability assays, both of which demonstrated tight junction formation and durable barrier function for up to 21 days (**Fig. 2c**). To determine the directionality of TLR2 activation, WT monolayers were stimulated apically or basolaterally with Pam3CSK4 (1 μg/ml, **Methods**). Apical stimulation selectively induced expression of canonical TLR2 target genes, including *Cxcl2* and *Tnfα*, within 4 hours, whereas basolateral stimulation failed to elicit a comparable response (**Fig. 2d-e**). No induction was observed in TLR2-deficient monolayers, confirming the specificity of apical TLR2 signaling (**Extended Data Fig. 2a,c**). Notably, TLR2-deficient monolayers also displayed impaired barrier integrity (**Extended Data Fig. 2b**), consistent with previous reports linking TLR2 to tight-junction regulation^40,41^. Bulk RNA-seq further showed strong induction of inflammatory cytokines and chemokines, including *Tnfα* and *Ccl20*, specifically in WT monolayers stimulated apically with Pam3CSK4, but not in unstimulated or TLR2 cKO cultures (**Fig. 2f**, **Supplementary Table 3**). Together, these findings indicate that stem-enriched epithelial monolayers respond selectively to apical TLR2 activation, supporting a model in which Lgr5^+^-ISCs can sense luminal microbial cues.

**Fig. 2.**
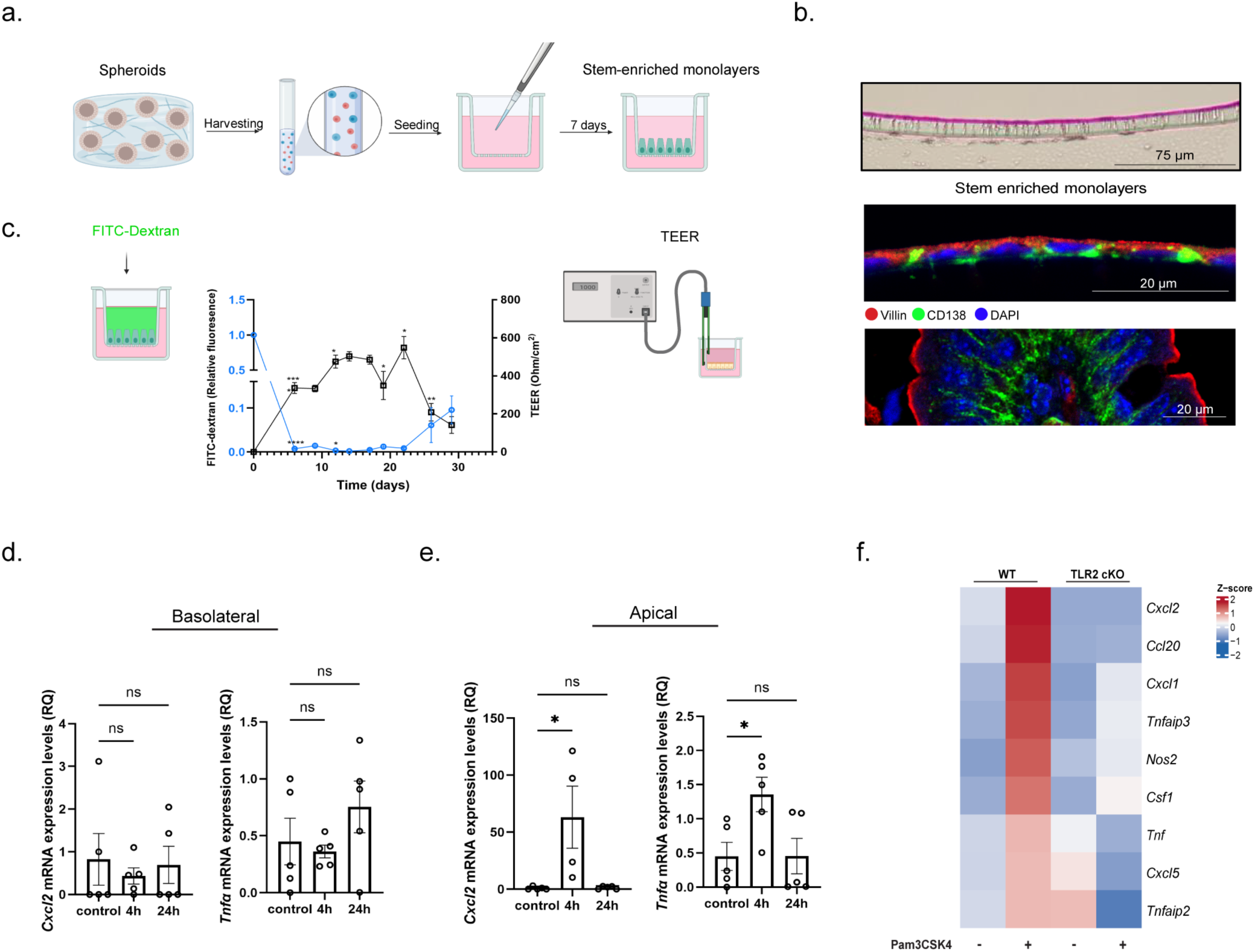
Primary ileal monolayers sense luminal content via TLR2. (**a**) Polarized monolayer generation from primary mouse SI spheroids scheme. (**b**) Representative images of primary mouse SI polarized monolayers using H&E staining (upper panel); scale bar, 75μm. IFA of primary monolayer (mid) or top of the villus (lower) stained with Villin (red), CD138 (green), and DAPI (blue); scale bar, 20μm. (**c**) Schematic representation of monolayer integrity assay, with FITC-dextran permeability quantification (left) or transepithelial electrical resistance (TEER) measurements (right) over 30 days of primary monolayer cultures. (**d-e**) Relative quantification (RQ) of TLR2 target genes, *Cxcl2* and *Tnfα* expression levels in stem-enriched monolayers after 4h or 24h of *ex vivo* apical (d) and basolateral (e) activation with TLR2/TLR1 agonist (Pam3Csk4, 1μg/mL) or without (control) using qPCR. Data are presented as mean ± SEM. *n*=5 biological repetitions per condition, *ns* (non-significant), \**p* <0.05, one-way ANOVA. (**f**) Heatmap of top TLR2 target genes from WT vs TLR2 cKO stem-enriched monolayers after 4h of *ex vivo* apical activation with or without Pam3Csk4 induction (1μg/mL), tested by bulk RNA-seq. Average relative expression (mean Z score of log_2_(TPM+1), color bar). *n*=3 vs 2 for WT or 3 vs 3 for TLR2 cKO biological repetitions for monolayers with or without Pam3Csk4 induction for 4h, respectively.

### Intestinal stem cell sensing via TLR2 is required for proper Paneth cell maturation

The ISC niche resides at the crypt base, a compartment generally protected from direct microbial exposure in a steady state^44,45^. Based on our observations that TLR2 is enriched in Lgr5^+^-ISCs and activated apically by MAMPs, we hypothesized that TLR2 may modulate ISC function by sensing microbial cues that reach this protected crypt environment *in vivo*. To test this, we examined the TLR2 cKO mouse model^42^, focusing on the ileum, where *Tlr2* expression is highest (**Fig. 1d**). Histological analysis of ileal sections from WT, TLR2 cKO, and TLR4 cKO mice revealed normal morphology in WT and TLR4 cKO animals, whereas TLR2 cKO mice showed marked bacterial accumulation and an increase in granular Paneth-like cells at the crypt base (**Fig. 3a**). In line with TLR4 being undetectably expressed in the SI in our scRNA-seq analysis (**Fig. 1a**), no noticeable phenotype was detected in TLR4 cKO. These findings suggested that TLR2 signaling in Lgr5^+^-ISCs may be required to maintain proper Paneth cell identity, and that its loss may lead to the accumulation of immature Paneth-lineage intermediate cells^43,44^. To further define the role of TLR2 in Lgr5^+^-ISCs, we crossed TLR2 cKO mice with Lgr5-2A-EGFP reporter mice^45^ to generate TLR2 cKO-Lgr5-GFP animals, enabling isolation of GFP^+^ Lgr5^+^-ISCs. Flow cytometric analysis revealed a marked reduction in GFP^high^ Lgr5^+^-ISCs and a milder reduction in GFP^low^ TA cells in TLR2 cKO mice (**Extended Data Fig. 3a-b**). Bulk RNA-seq of 5,000 sorted Lgr5^+^-ISCs from WT and TLR2 cKO mice identified extensive transcriptional changes, with 539 upregulated and 598 downregulated differentially expressed genes (DEGs) in the absence of TLR2 (**Fig. 3b**, **Supplementary Table 4** and **Methods**). Notably, among the most significantly reduced genes were those associated with defense responses to bacteria and AMPs, key innate effectors of epithelial defense ^46,47^. Gene Ontology analysis confirmed that antibacterial defense pathways were among the most significantly downregulated biological processes (**Fig. 3c**, **Supplementary Table 5**). In particular, members of the Defensin and Reg3 AMP families, typically expressed by Paneth cells and enterocytes, respectively, were reduced in TLR2-deficient Lgr5^+^-ISCs (**Fig. 3d**, **Supplementary Table 4**), suggesting that TLR2 signaling in GFP^high^ cells (mostly Lgr5^+^-ISCs) contributes to epithelial antimicrobial programming.

**Fig. 3.**
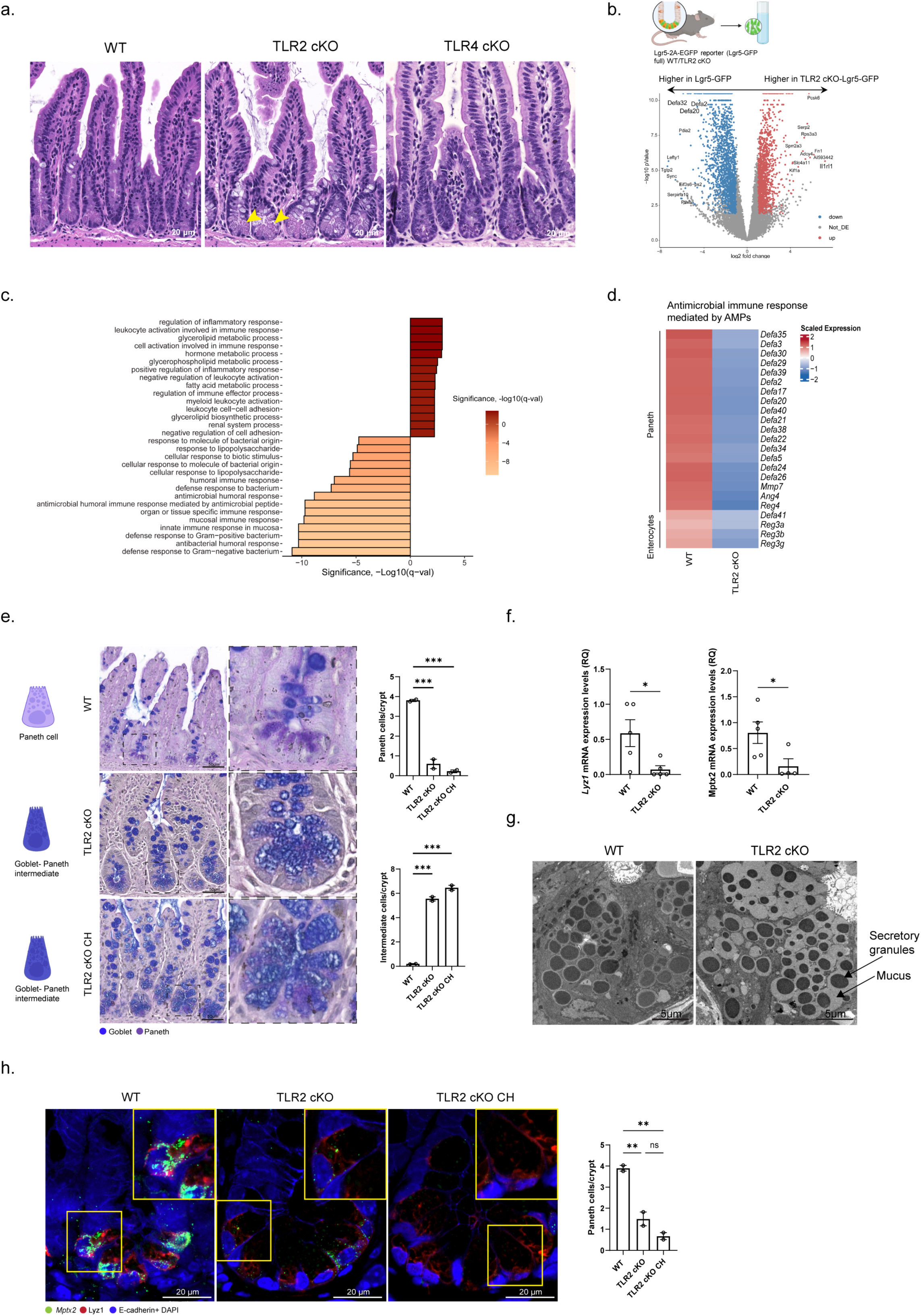
Disruption of Paneth cell maturation in TLR2 cKO. (**a**) Representative H&E images of SI ileum sections from WT, TLR2 cKO, or TLR4 cKO mice; scale bars, 20μm; yellow arrows show the changes in granular cells at the bottom of the crypt in TLR2 cKO mice. (**b-d**) Comparison of Lgr5^+^-ISCs from WT or TLR2 cKO mice using bulk-RNA sequencing. (b) Experimental scheme for GFP-labeled Lgr5^+^-ISC sort (top) and volcano plot showing log2fc estimates and -log(FDR q-value) of top ten upregulated (right) and downregulated (left) differentially expressed genes (DEG) in Lgr5^+^-ISCs of TLR2 cKO (*n*=2) compared to WT (*n*=3) mice. Colored points correspond to padj ≤ 0.05 and |log2FoldChange| ≥ 1. Color coding is shown in the bottom-right corner. (c) Gene ontogeny (GO) biological processes showing hypergeometric enrichment of significantly upregulated and downregulated DEGs in Lgr5+-ISCs of TLR2 cKO (*n*=2) vs. WT (*n*=3) mice. Plotted is the negative log10 of enrichment padj. The direction of enrichment is shown as positive for biological processes enriched in TLR2 cKO, and negative for those enriched in WT. (d) Heatmap of the top DE antimicrobial peptides from WT (*n*=3) vs. TLR2 cKO (*n*=2) mice. Average relative expression (mean Z score of log_2_(TPM+1), color bar). (**e-g**) Goblet-Paneth intermediate cells accumulation is observed in TLR2 cKO. (e) Representative images of Periodic-Acid Schiff (PAS) and Alcian Blue (AB) staining of ileal sections from WT (top), TLR2 cKO (mid), and TLR2 cKO chimeric (CH, bottom) mice. The number of Paneth cells and goblet-Paneth intermediate cells per crypt was quantified (right); *n* = 10 fields per mouse, 2 mice per group; data are mean ± SEM; \*\*\**p* < 0.001, one-way ANOVA. scale bar, 50μm. Inset, x4 magnification. (f) RQ of *Lyz1* and *Mptx2* expression levels in WT vs. TLR2 cKO using qPCR. Data are presented as mean ± SEM. *n*=5 WT vs 4 of TLR4 cKO mice, \**p* <0.05, two-tailed *Student’s t-test*. (g) Transmission electron microscopy (TEM) of Paneth cell eosinophilic granules in SI ileal segments of WT (left) and TLR2 cKO (right) mice; Scale bar, 5μm. (h) Mature Paneth cells staining in SI ileal crypts of WT (left), TLR2 cKO (mid), or TLR2 cKO chimera (CH, right). Single-molecule fluorescence *in situ* hybridization (smFISH) of *Mptx2* (green) co-stained with Lyz1 antibody (red). E-Cadherin stains IEC boundaries, while DAPI stains the nucleus (blue). Scale bar, 20μm. Insets, x1.4 magnification. Lyz1+ *Mptx2*+ Paneth cells per crypt were quantified (right); *n*=10 fields per mouse, 2 mice per group. Data are presented as mean ± SEM. *ns*, non-significant, \*\**p*<0.01, one-way ANOVA.

To link AMP loss with the accumulation of Paneth-like cells in TLR2 cKO crypts, we stained ileal tissue with Alcian Blue-Periodic Acid-Schiff (AB-PAS, **Methods**), which distinguishes goblet and Paneth lineages by acidic and neutral mucins^48,49^. TLR2 cKO crypts contained cells with goblet-Paneth intermediate features, indicating impaired terminal maturation of the Paneth lineage^43,50^ (**Fig. 3e**). Consistent with this, qPCR revealed reduced expression of mature Paneth cell markers and AMP genes, including *Lyz1* and *Mptx2* (**Fig. 3f**). Transmission electron microscopy (TEM) further revealed smaller, structurally altered eosinophilic granules, along with intracellular mucin accumulation, supporting the presence of immature secretory intermediate cells in TLR2 cKO crypts (**Fig. 3g**). To determine whether this phenotype reflected epithelial rather than hematopoietic TLR2 deficiency, we generated bone marrow chimeras in which irradiated TLR2 cKO mice were reconstituted with WT GFP-labeled immune cells (**Extended Data Fig. 3c**). Successful immune reconstitution was confirmed by the presence of GFP^+^ immune cells in the lamina propria (**Extended Data Fig. 3d-e**). Despite restoration of a TLR2-sufficient immune compartment^51^, chimeric TLR2 cKO mice still accumulated goblet-Paneth intermediate cells (**Fig. 3e**) and showed markedly reduced expression of Lyz1 and Mptx2 by immunofluorescence and smFISH (**Fig. 3h** and **Extended Data Fig. 3f**). Together, these data demonstrate that TLR2 expression in Lgr5^+^-ISCs is required for proper secretory lineage differentiation, particularly for mature Paneth cell formation and the establishment of the Paneth-specific AMP program.

### TLR2 regulates Lgr5^+^-ISC-dependent antimicrobial programs across epithelial lineages

Given the defects in Paneth maturation and reduced AMP expression observed in TLR2 cKO mice, we next examined the broader epithelial consequences of TLR2 loss at single-cell resolution. We performed scRNA-seq on epithelial cells isolated from the distal SI of chimeric WT and TLR2 cKO mice (**Methods**). Unsupervised clustering identified 12 epithelial subpopulations based on established lineage markers^4,48^ (**Fig. 4a, Extended Data Fig. 4a**, and **Supplementary Table 6**). TLR2 loss induced widespread changes in epithelial composition. Chimeric TLR2 cKO mice showed a marked reduction in stem and TA populations (**Fig. 4a**), consistent with the depletion observed in TLR2-deficient Lgr5-GFP mice (**Extended Data Fig. 3a-b**). This decrease correlated with reduced epithelial proliferation, as shown by cell-cycle scoring and Ki67 staining (**Fig. 4b** and **Extended Data Fig. 4b,d**), and not with an increase in apoptosis (**Extended Data Fig. 4e**). In parallel, ISC transcriptional changes included increased expression of genes associated with oxidative stress, such as Gpx2 and G6pdx, as well as immune surveillance pathways, including MHC class II antigen presentation, which may suggest a compensatory response in ISCs^3,15,52^ (**Extended Data Fig. 4c**, **Supplementary Table 7**). Reduction of stem and TA populations was accompanied by a relative expansion of differentiated lineages, particularly mature enterocytes and tuft cells (**Fig. 4a,c**). Pseudotime analysis reconstructed three major epithelial trajectories and showed that TLR2-deficient cells were shifted toward more advanced differentiation states along both the enterocyte and tuft lineages (**Fig. 4d** and **Extended Data Fig. 4f**). Along the enterocyte trajectory, TLR2-deficient cells progressed further toward terminal villus-tip identity, characterized by the expression of apolipoproteins such as Apoa4^53^ (**Fig. 4e**, **Supplementary Table 7**). Consistently, analysis of enterocyte zonation scores^53^ revealed altered spatial positioning already within the mature enterocyte compartment (**Extended Data Fig. 4g**).

**Fig. 4.**
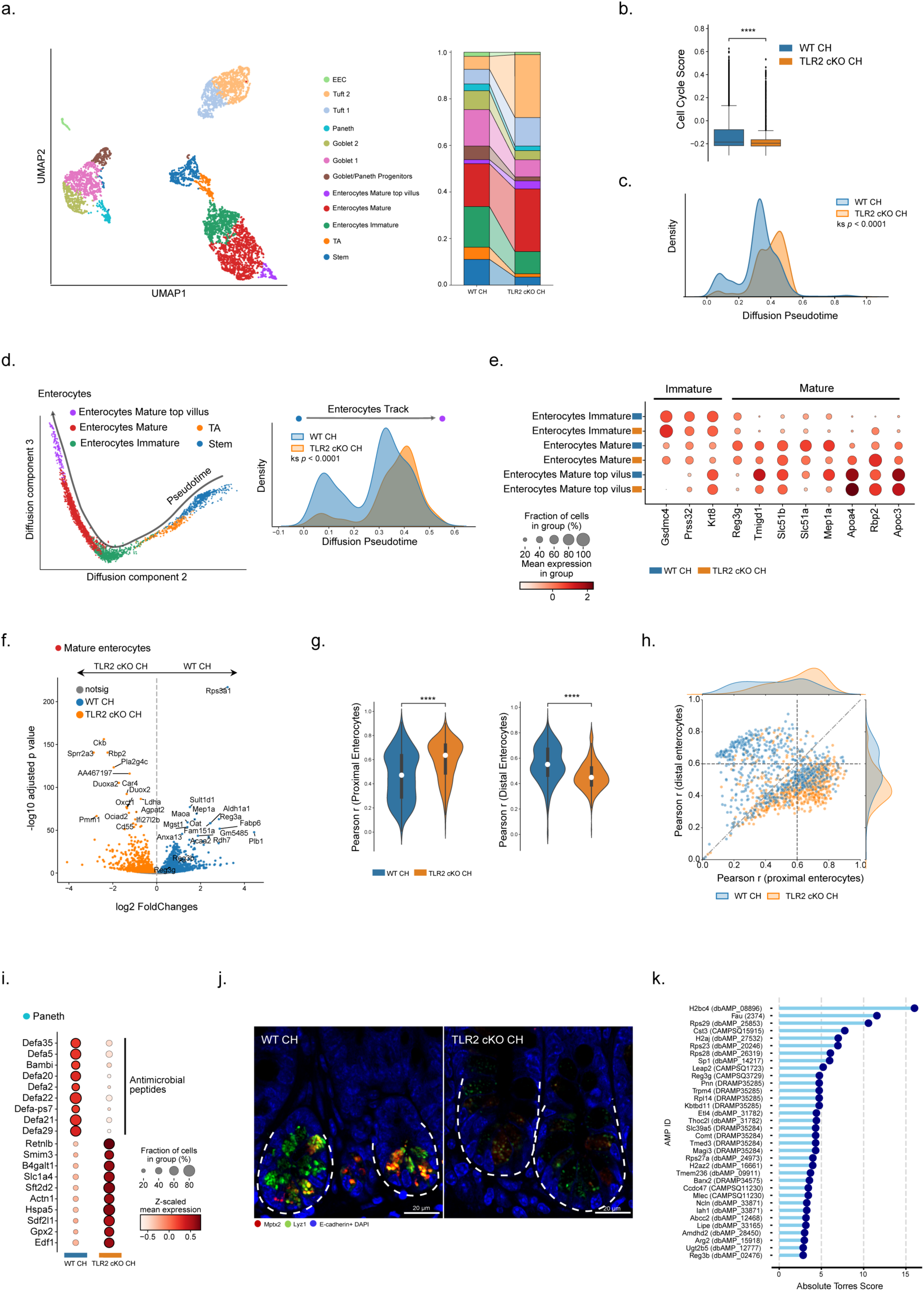
Reduction of the AMP program in the TLR2 cKO chimera at single cell resolution. (**a**) Two-dimensional Uniform Manifold Approximation and Projection (UMAP) graphical representation of EpCAM^+^ single cell RNA-sequencing (scRNA-seq), colored by graph clustering per different clusters from chimeric (CH) WT and TLR2-KO mice (left) and stacked bar of relative cell subset abundance in chimeric (CH) WT and TLR2 cKO (right). Data are presented as mean ± SEM. *n*=2 mice per condition. TA, transient amplifying cells; EEC, enteroendocrine. (**b**) Box-and-whisker plot of cell cycle signature score in chimeric (CH) WT vs TLR2 cKO. *n*=2 mice per condition. Boxes represent median and Interquartile Range (IQR); whiskers extend to 1.5× IQR. Mann-Whitney U test; \*\*\*\**p*<0.0001. (**c**) Diffusion pseudotime (dpt) score comparing the differentiation state of chimeric (CH) WT (blue) and TLR2 cKO (orange). Dpt scores showing a right-shift peak in TLR2 cKO cells, indicating accelerated differentiation compared to WT. (**d**) Diffusion map of enterocytes differentiation trajectory colored by cell subset in chimeric (CH) WT and TLR2 cKO (left) and diffusion pseudotime (dpt) score comparing the enterocytes differentiation state of chimeric (CH) WT (blue) and TLR2 cKO (orange) (right). Dpt scores showing a right-shift peak in TLR2 cKO cells, indicating accelerated differentiation compared to WT. (**e**) Dot plot of immature and mature enterocyte cell markers comparing enterocyte subsets of chimeric (CH) WT (blue) vs. TLR2 cKO (orange) mice. The size of the dot indicates the proportion of cells expressing a gene, while the color indicates the mean expression level. (**f**) Volcano plot showing log2fc estimates and the negative log10 of enrichment padj. of top upregulated (right) and downregulated (left) DEG in mature enterocytes of chimeric (CH) WT (*n*=2) compared to TLR2 cKO (*n*=2) chimeras. Colored points correspond to padj ≤ 0.05 and |log2FoldChange| ≥ 1. Color coding is shown on the top left. (**g**) Distribution of proximal (left) and distal (right) enterocyte signatures in mature enterocytes of chimeric (CH) WT (blue) vs. TLR2 cKO (orange) mice; Significance was determined by Pearson’s *r* test, \*\*\*\**p*<0.0001. (**h**) Scatter plot comparing Pearson correlation coefficients (r) of proximal and distal enterocyte gene signatures in chimeric (CH) WT and TLR2 cKO mature enterocyte cells. The x-axis represents the Pearson r values for proximal enterocyte signature genes, and the y-axis shows Pearson r values for distal enterocyte signature genes. Each point corresponds to a cell within the respective signature. The colors represent different conditions: blue for WT and orange for TLR2 cKO. (**i-j**) Antimicrobial peptides expression in Paneth cells. (i) Dot plot of top DEG in Paneth cells from chimeric (CH) WT vs. TLR2 cKO mice. The size of the dot indicates the proportion of cells expressing a gene, while the color indicates the average relative expression (mean Z score of log2(TPM+1)). (j) Reduction of AMP expressed by Paneth cells in TLR2 cKO. IFA of Lyz1 (green), Mptx2 (red), E-Cadherin, and DAPI co-staining in chimeric (CH) WT (left) and TLR2 cKO (right) mice. **(k)** Uncharacterized top AMP-encrypted candidates in mature enterocytes derived from DEG upregulated in chimeric WT compared to TLR2 cKO. X-axis: gene list, Y-axis: absolute Torres score.

Because enterocyte function varies along the SI axis^4,18^, we next asked whether TLR2 loss altered regional enterocyte identity. Mature enterocytes from TLR2-deficient ileum downregulated distal markers such as *Fabp6* and *Plb1*, involved in lipid metabolism, while upregulating proximal markers including *Ckb* and *Rbp2* (**Fig. 4f**, **Supplementary Table 7**). Correlation with a publicly available proximal and distal enterocyte reference dataset ^4,56^ confirmed this shift toward proximal-like identity (**Fig. 4g,h**). Importantly, the enterocyte AMP program, which is enriched in the distal SI and dominated by Reg3 family genes^4,54^, was also reduced in TLR2-deficient enterocytes (**Fig. 4e,f** and **Supplementary Table 7**).

Analysis of Paneth cells at single cell resolution similarly revealed reduced antibacterial AMP signatures in chimeric TLR2 cKO mice (**Extended Data Fig. 4h**). Canonical AMP genes, including multiple Defensins, were markedly downregulated (**Fig. 4i**, **Supplementary Table 7**), and reduced Lyz1 and Mptx2 staining further confirmed loss of Paneth cell antimicrobial output (**Fig. 4j**). Of note, in addition to altered enterocyte identity, Paneth cells from chimeric TLR2 cKO ileum also shifted toward a more proximal-like state (**Extended Data Fig. 4j**), indicating that regional epithelial identity and antimicrobial output are both shaped by TLR2-mediated microbial sensing in ISCs.

To identify new TLR2-dependent AMP candidates in enterocytes and Paneth cells, we expand this analysis. We interrogated downregulated genes in enterocytes and Paneth cells in the TLR2 cKO for potential AMP properties^55,56^. While several known Defensin family members were identified in Paneth cells (**Extended Data Fig. 4i**, **Supplementary Table 8**), we additionally uncovered 34 candidate AMP-related, TLR2-dependent genes in mature enterocytes. Although the known *Reg3b* and *Reg3g* were included in our list, the strongest candidates were *H2bc4*, *Fau*, *Rps29*, and *Cst3*, all of which were enriched in TLR2-proficient enterocytes (**Fig. 4k**, **Supplementary Table 8**). Antimicrobial activity has previously been reported for these proteins^55,57–59^, suggesting that the ileal enterocyte AMP repertoire is broader than previously appreciated and is regulated, at least in part, through TLR2 activation in Lgr5^+^-ISCs.

### Epithelial and ISC-specific deletion of TLR2 disrupts epithelial antimicrobial programs

To define the epithelial-intrinsic role of TLR2, we generated a constitutive epithelial-specific knockout driven by Villin-Cre (TLR2^ΔIEC^; **Fig. 5a** and **Extended Data Fig. 5b**). Similar to chimeric TLR2 cKO mice, TLR2^ΔIEC^ animals displayed impaired Paneth cell maturation and accumulation of goblet-Paneth intermediate cells in ileal crypts (**Fig. 5b** and **Extended Data Fig. 5a,c**). Paneth-specific AMP expression was also significantly reduced, as shown by diminished Lyz1 and Mptx2 staining (**Fig. 5c**). To directly assess the role of TLR2 in Lgr5^+^-ISCs and their progeny, we generated a lineage-tracing model by crossing Lgr5-CreER^T2^-TLR2^fl/fl^ to Rosa26^LSL-tdTomato^ mice (TLR2^ΔISC+tdTom^; **Methods**, **Fig. 5d**). 48 hours after tamoxifen induction, flow cytometric analysis revealed a reduction in tdTomato^+^ epithelial progeny in TLR2^ΔISC+tdTom^ mice relative to controls (**Fig. 5e**). Consistent with our scRNA-seq findings, tissue analysis showed reduced numbers of tdTomato^+^ cells and decreased proliferation in the crypt zone (**Fig. 5f**). TUNEL staining showed no increase in cell death among tdTomato^+^ cells in TLR2-deficient crypts (**Extended Data Fig. 5d**), indicating that the reduction reflected impaired proliferation rather than increased apoptosis. At 5 days after induction, TLR2-deficient progeny showed reduced *Tlr2* expression and decreased proliferation (**Extended Data Fig. 5e,f**). Importantly, TLR2-deficient Paneth progeny (tdTomato^+^ Lyz1^+^) exhibited a significant reduction in the Paneth marker Lyz1 (**Fig. 5g**). Together, these findings indicate that TLR2-sensing by Lgr5^+^-ISCs serves not only to maintain the Lgr5^+^-ISC niche but also as a key regulator of the epithelial antimicrobial program.

**Fig. 5.**
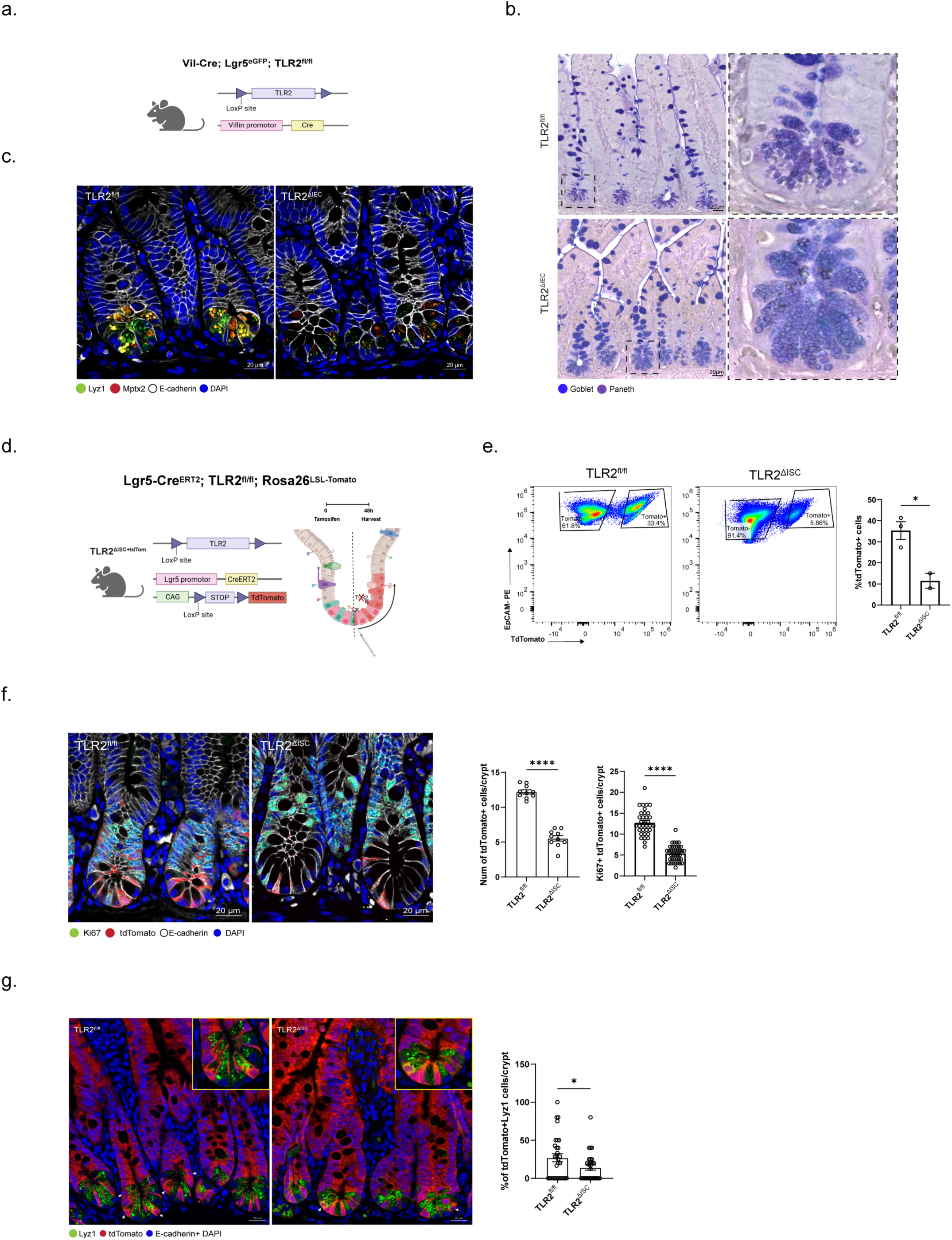
Characterization of epithelial and ISC-specific TLR2 deletion. (**a-c**) Epithelial-specific TLR2 deletion (TLR2^ΔIEC^) mouse model. (a) Experimental scheme for the generation of epithelial-specific TLR2 deletion by crossing Vil-Cre to TLR2^fl/fl^ mouse model (TLR2^ΔIEC^). (b) Representative images of PAS and AB staining of ileum sections from TLR2^fl/fl^ (upper panel) and TLR2^ΔIEC^ (lower panel). scale bar: 20 μm. Insets, x5 magnification. (c) AMP (Lyz1 and Mptx2) expressed by Paneth cells staining in TLR2^fl/fl^ (left) and TLR2^ΔIEC^ (right) ileal sections. IFA of Lyz1 (green), Mptx2 (red), E-cadherin (white) and DAPI. E-cadherin (white) stains IEC boundaries; scale bar: 20 μm. (**d-g**) Lgr5+-ISC-specific TLR2 deletion (TLR2^ΔISC^) mouse model. (d) Schematic representation of Lgr5+-ISC-specific TLR2 deletion lineage tracing (TLR2^DISC+tdTom^) generation by crossing Lgr5+-ISC-CreER^T2^-TLR2^fl/fl^ mice to Rosa26-tdTomato mice. TdTomato cells are progeny of TLR2-deleted Lgr5+-ISC after one injection of Tamoxifen (2mg) at 48h (d-f) or 5 days (g). (e) Representative flow cytometry plot of the percentage of TdTomato^+^ EpCAM^+^ cells among EpCAM^+^ cells in TLR2^fl/fl^ vs. TLR2^ΔISC^ mice (left panel); quantification of TdTomato^+^ EpCAM^+^ percentages in both groups is depicted on a bar graph (right panel); *n*=3 mice per TLR2^fl/fl^ and 2 mice per TLR2^ΔISC^. Data are presented as mean ± SEM. \**p*<0.05, two-tailed *Student’s t-test*. (f) IFA of tdTomato (red), Ki67 (green), E-cadherin (white), and DAPI of distal SI, 48h after Tamoxifen induction. E-cadherin (white) stains IEC boundaries; scale bar, 20 μm. (left panel). Quantification of tdTom+ cells (mid) and of Ki67+ tdTom+ cells (Right) per crypt. Data are presented as mean ± SEM. 10 fields (tdTom+) or 20 fields (tdTom+ Ki67+) per mouse. \*\*\*\**p* < 0.001, two-tailed *Student’s t-test*. (g) IFA of tdTomato (red), Lyz1 (green), E-cadherin+DAPI (blue) of distal SI, 5 days after Tamoxifen induction; scale bar, 20 μm. (left panel). Quantification of tdTom+ Lyz1+ cells (right panel) per crypt. Data are presented as mean ± SEM. 20 fields per mouse and *n*=2 mice. \**p* < 0.05, two-tailed *Student’s t-test*.

### TLR2 deficiency induces expansion of bacterial-sensing tuft cells

Reduction in AMP programs upon TLR2 depletion compromises a major arm of SI antimicrobial defense, raising the possibility that compensatory epithelial programs are activated. Tuft cells are specialized chemosensory epithelial cells implicated in type 2 immunity and intestinal homeostasis^5–7^. More recently, tuft cells have also been implicated in bacterial sensing through receptors such as the succinate receptor SUCNR1^60–62^ and the vomeronasal receptor Vmn2r26, which detects the bacterial metabolite N-undecanoylglycine (N-C11-G) and promotes prostaglandin D2-mediated antibacterial responses^63^.

In TLR2-deficient mice, flow cytometry revealed a significant expansion of tuft cells (SSC^low^ CD24^high^), accompanied by increased *Dclk1* expression in sorted ileal EpCAM^+^ cells (**Extended Data Fig. 6a,b**). Bulk RNA-seq further showed upregulation of markers associated with both Tuft-1 and Tuft-2 subsets^12,66^ (**Extended Data Fig. 6c**, **Supplementary Table 4**). Since tuft differentiation has recently been described as a linear progression from progenitor to Tuft-1 to Tuft-2 states^64^, we examined these populations by histology. Immunofluorescence analysis confirmed expansion of both DCLK1^+^CD45^-^ Tuft-1 and DCLK1^+^CD45^+^ Tuft-2 subsets^68–71^ in TLR2-deficient ileum (**Extended Data Fig. 6d,e**). This increase was consistent with our scRNA-seq data, which also showed preferential enrichment of Tuft-1 and Tuft-2 populations in chimeric TLR2 cKO mice (**Fig. 4a**). Pseudotime analysis further revealed a shift toward terminal tuft differentiation in TLR2-deficient epithelial cells (**Extended Data Fig. 6f**). Notably, tuft cells from TLR2-deficient mice showed elevated expression of the Tuft-2-associated bacterial-sensing receptors *Sucnr1* and *Vmn2r26* (**Extended Data Fig. 6g**, **Supplementary Table 9**). These findings suggest that expansion of bacterial-sensing Tuft-2 cells may represent a compensatory mechanism that could support antimicrobial defense and host-microbial symbiosis when TLR2 signaling is lost.

### Stem cell sensing of gut bacteria induces Paneth cell differentiation

The SI contains a substantial microbial load, increasing from approximately 10^4^–10^5^ CFU/ml in the duodenum to 10^7^–10^8^ CFU/ml in the distal ileum^12,65^. Because our data indicated that Lgr5^+^-ISCs sense microbial signals through apical TLR2 and initiate antimicrobial programs, we next tested the contribution of the microbiota itself using germ-free (GF) mice. Flow cytometric analysis showed a significant reduction in Lyz1^+^ Paneth cells in GF mice relative to specific pathogen-free (SPF) controls (**Fig. 6a**), which was confirmed by AB-PAS staining showing a marked decrease in mature Paneth cells in GF mice (**Fig. 6b**). Consistently, expression of the Paneth AMP marker *Mptx2* and the enterocyte-enriched AMP *Reg3g* was reduced in GF ileum (**Fig. 6c**). Notably, *Lgr5* and TLR2-related target genes were likewise downregulated in GF intestinal epithelium (**Extended Data Fig. 7a**), supporting a role for microbiota-derived signals in ISC function. To determine whether microbial exposure could restore AMP expression and Paneth cell differentiation, we colonized GF mice with microbiota isolated from SPF WT donors (**Extended Data Fig. 7b**). After 14 days, colonized mice exhibited Paneth cell differentiation by AB-PAS staining (**Fig. 6d**), and flow cytometry showed a trend toward increased Lyz1+ epithelial cells (**Extended Data Fig. 7c**). These data support a model in which microbiota-derived signals are associated with Paneth differentiation, potentially through the TLR2 pathway under homeostatic conditions.

**Fig. 6.**
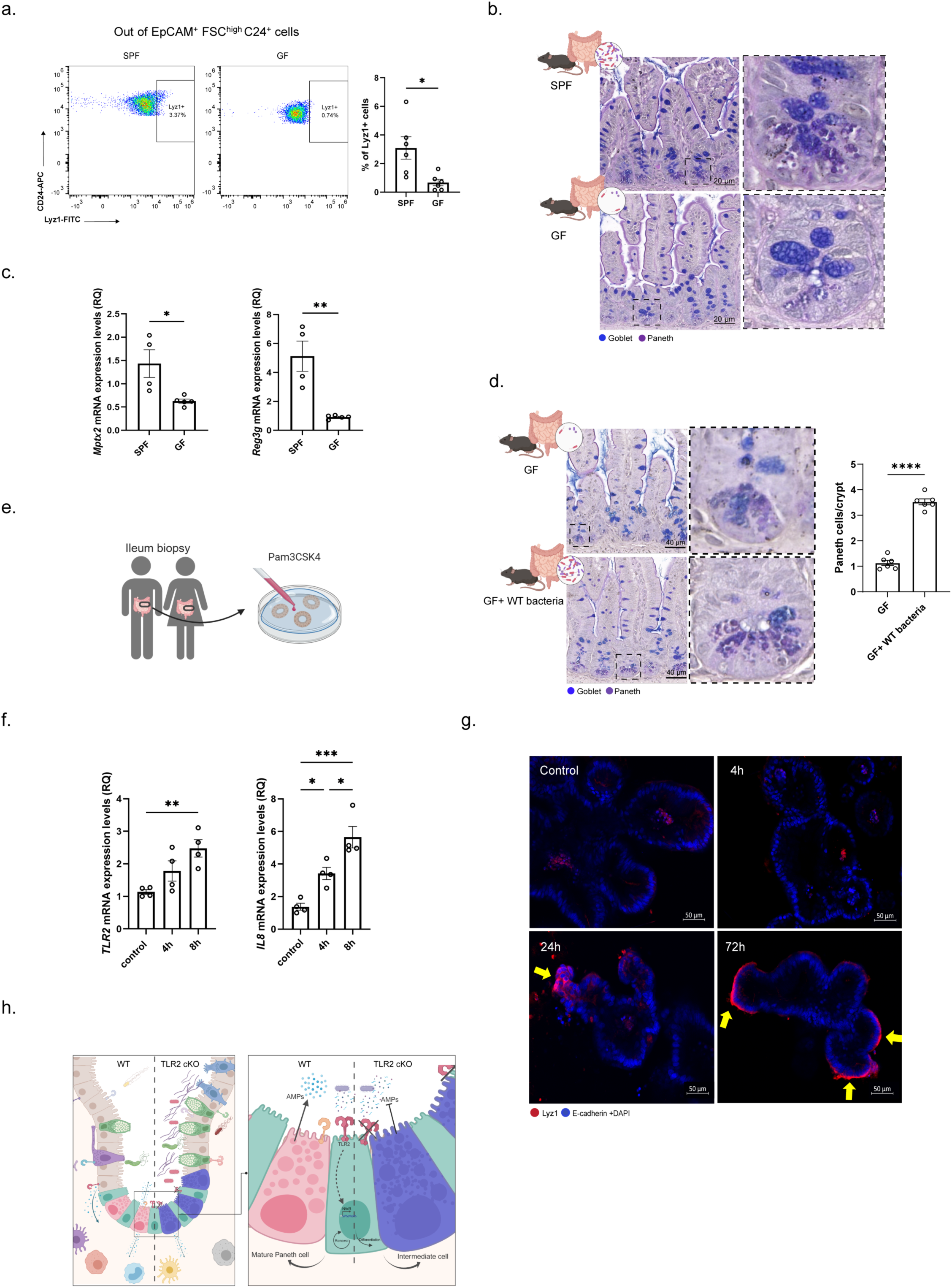
TLR2 activation by microbiota induces Paneth differentiation. (**a-d**) The involvement of microbiota in TLR2 activation and Paneth differentiation. (a) Representative flow cytometry plot (left) and quantification (right) of the percentage of Lyz1^+^ cells among EpCAM^+^ cells in ileal crypts of SPF vs. GF mice. *n*=6 mice per group; data are presented as mean ± SEM. \**p*< 0.05, two-tailed *Student’s t-test*. (b) Representative images of PAS and AB staining of ileum sections from SPF (upper panel) and GF (lower panel) mice; Scale bar, 20μm. Inset, x5 magnification. (c) Relative quantification (RQ) of *Mptx2 and Reg3γ* expression levels, key Paneth and enterocyte AMPs, in EpCAM^+^ cells of GF (*n*=5) vs. SPF (*n*=4) mice using qPCR. Data are presented as mean ± SEM; *ns* (non-significant), \**p*<0.05, \*\**p*<0.01, two-tailed *Student’s t-test*. (d) GF colonization experiment. Representative images of PAS and AB staining of ileum sections (Left) from GF (upper panel) or GF colonized with WT bacteria (lower panel); *n*=4 mice per group, scale bar, 20μm. Inset, x5 magnification. Quantification of Paneth cells per crypt of GF vs GF colonized with WT bacteria (right). 6 fields of view per condition; data are presented as mean ± SEM. \*\*\*\**p*< 0.0001, two-tailed *Student’s t-test*. (**e-g**) Human ileal spheroids *in vitro* activation assay. (e) Experimental scheme of generating spheroids from human ileum biopsies and activation with TLR2/TLR1 agonist (Pam3Csk4). (f) Relative quantification (RQ) of *TLR2* and *IL8* expression levels in spheroids generated from ileal biopsies after 4h or 8h of *ex vivo* activation with TLR2/TLR1 agonist (Pam3Csk4, 1μg/mL) or without (control) using qPCR; *n*=4 biological repetitions per group. Data are presented as mean ± SEM. *ns* (non-significant), \**p*<0.05, \*\**p*<0.01, \*\*\**p* <0.001, one-way ANOVA. (g) Human Paneth cell expansion following Pam3Csk4 induction. IFA of LYZ1 (red), E-cadherin (blue), and DAPI in fixed differentiated spheroids after 4h, 24h, and 72h of *ex vivo* activation with TLR2/TLR1 agonist (Pam3Csk4, 1μg/mL) or without (control); scale bar, 50μm. Yellow arrows, LYZ1^+^ crypts. (**h**) Suggested model for the symbiotic relationship of the Lgr5+-ISC with microbiota mediated by TLR2 signaling.

Impaired AMP production is associated with inflammatory bowel disease, such as Crohn’s disease, where reduced ileal α-defensin expression has been observed^66–69^. We therefore asked whether TLR2 activation could similarly promote Paneth cell differentiation in human organoids. As previously shown, goblet and Paneth lineages in murine organoids remain undifferentiated, often co-expressing markers of both cell types^15,50,70^. This intermediate state was recapitulated *in vivo* in constitutive or epithelial TLR2 KO mice, indicating that TLR2 activation in Lgr5^+^-ISC is required for Paneth lineage commitment, in addition to the Wnt pathway activation shown before^71,72^. In human SI organoid cultures, secretory cell differentiation is limited, and mature Paneth cells are particularly rare^50,73,74^. We first reanalyzed public human ileal scRNA-seq data^75^ and found enriched expression of TLR1 and TLR2 in ISC and TA populations (**Extended Data Fig. 7d**). We then cultured human ileal-derived stem-enriched spheroids and stimulated them with Pam3CSK4 for 4 or 8 hours (**Methods, Fig. 6e**). TLR2 activation induced expression of *TLR2* itself and the downstream target gene, *IL8* (**Fig. 6f**). To assess whether TLR2 activation promotes human Paneth differentiation, we treated human-seeded ileal organoids with Pam3CSK4 (1μg/ml) for 4, 24, or 72 hours. Prolonged stimulation induced progressive accumulation of LYZ1 protein at the base of crypt-like domains, particularly after 72 hours (**Fig. 6g**). These results demonstrate that human ISCs are functionally responsive to TLR2 activation and that TLR2 signaling promotes human Paneth cell differentiation.

Together, these findings support a model in which TLR2^+^ ISCs act as microbial sensors, integrating luminal microbial signals through apical TLR2 activation to regulate stem cell maintenance, epithelial AMP programs, and Paneth cell maturation (**Fig. 6h**). In the absence of TLR2, this feedback loop is disrupted, resulting in reduced ISC numbers, impaired Paneth lineage commitment, accumulation of goblet-Paneth intermediate cells, diminished AMP secretion, and potential dysbiosis.

## Discussion

In this study, we identify TLR2 as an epithelial-intrinsic sensor enriched in Lgr5^+^-ISCs that enables luminal microbial sensing required for epithelial homeostasis and differentiation at steady state. We show that TLR2-mediated sensing in Lgr5^+^-ISCs induces an epithelial antimicrobial program and promotes lineage specification toward Paneth and enterocyte states enriched for AMP expression, particularly in the distal small intestine (SI). Using constitutive, epithelial-specific, and Lgr5^+^-ISC-specific TLR2 deletion models, together with *ex vivo* mouse and human organoid systems, we demonstrate that epithelial-intrinsic TLR2 signaling is required to maintain the ISC compartment and epithelial innate immune function. Loss of TLR2 impairs Paneth cell differentiation, leads to the accumulation of goblet-Paneth intermediate cells, and reduces AMP expression across epithelial lineages. Moreover, Lgr5^+^-ISC fate-mapping experiments support a direct role for TLR2 in ISC regulation. Together, these findings position Lgr5^+^-ISCs not simply as passive recipients of niche-derived cues, but as active sensors of microbial signals that couple epithelial regeneration to host defense.

Previous studies have established that epithelial TLR2 contributes to mucosal integrity by regulating tight junctions^76^. Moreover, commensal sensing via TLR-MYD88 signaling has been linked to AMP induction, mainly in differentiated epithelial cells, particularly Paneth cells or mature enterocytes^54,77,78^. In parallel, mapping of TLR expression suggested that TLR5, rather than TLR2, is selectively enriched in Paneth cells^24^. Our findings expand this framework by identifying TLR2 as an ISC-enriched PRR and demonstrating that its activation in Lgr5^+^-ISCs can drive transcriptional and differentiation programs that shape epithelial antimicrobial defense. These data fill an important gap in understanding how stem cells integrate microbial information to regulate lineage allocation and tissue function.

A central conceptual advance of this study is the identification of apical TLR2 activation as a mechanism that may enable luminal sensing by adult stem cells. Although the intestinal crypt is generally considered a protected niche^81^, our monolayer experiments indicate that ISCs can detect luminal microbial-derived ligands via TLR2 signaling. This is particularly relevant in the ileum, where microbial burden is high^31,65^ and *Tlr2* expression is enriched. We propose that luminal sensing by ISCs provides a mechanism by which microbial abundance and composition can be translated into rapid changes in epithelial differentiation and antimicrobial output. Supporting this, colonization experiments in GF mice further confirm that microbial sensing is necessary for ISC renewal and differentiation. Thus, we propose that Lgr5^+^-ISCs contribute to maintaining a symbiotic relationship with the gut microbiota via apical TLR2 activation, which ensures, on one hand, a proper epithelial renewal and, on the other hand, bacterial defense mechanisms. More broadly, these results support the idea that adult stem cells in barrier tissues may function as environmental sensors, integrating local cues to coordinate tissue adaptation.

Our data further demonstrate that TLR2 signaling is required for proper Paneth cell maturation. In the absence of TLR2, ileal crypts accumulate immature goblet-Paneth intermediate cells, with reduced expression of mature Paneth markers and disrupted granule architecture. This phenotype is consistent with a block in terminal secretory differentiation rather than simple lineage loss. These findings align with prior studies showing that Paneth cell maturation depends on inflammatory signaling inputs, including IFNγ, IL-17, and IL-22^15,74,79–81^. Our results add microbial sensing via TLR2 in ISC to support Paneth cell maturation. This phenotype is consistent with findings by Brischetto et al., who showed that disruption of epithelial NF-κB signaling led to a similar expansion of the immature secretory intermediate^82^. Given that TLR2 signals through the MYD88-NF-κB axis^51^, loss of TLR2 may impair the transcriptional program required for Paneth differentiation and AMP production.

Beyond Paneth cells, TLR2 loss reshaped epithelial organization more broadly. Single cell analysis revealed reduced stem and transit-amplifying populations, decreased proliferation, and a shift toward differentiated enterocyte and tuft lineages. In enterocytes, TLR2 deficiency was associated with loss of distal identity and reduced expression of AMPs, such as Reg3 family members. These results suggest that microbial sensing by ISCs contributes not only to lineage balance but also to regional epithelial specialization in the ileum. This is particularly important because the distal SI is a site of high microbial exposure and requires robust antimicrobial regulation^4,31^.

As antibiotic resistance becomes a real threat, the search for alternatives, such as natural AMPs, is on the rise^83–85^. The regulation of the known epithelial AMP levels by TLR2 expression led us to search for additional uncharacterized epithelial-AMP candidates. Our findings support that the epithelial AMP repertoire may be substantially broader than previously appreciated and is regulated, at least in part, by TLR2 signaling. In addition to reduced expression of canonical Paneth-derived Defensins, we identified a broader set of candidate AMP-related genes in enterocytes, including *H2bc4*, *Fau*, *Rps29*, and *Cst3*. Although the antimicrobial activity of some of these proteins has been described in other contexts^55,57–59^, their role in the intestinal epithelium remains largely unexplored. These results raise the possibility that enterocytes harbor an expanded antimicrobial arsenal tuned by stem-cell-derived microbial sensing. Functional studies will be needed to determine how these candidate peptides contribute to host defense, microbiota composition, and epithelial resilience.

Loss of TLR2 in Lgr5^+^-ISCs also triggered expansion of tuft cells, particularly the Tuft-2 subset associated with bacterial sensing^60–63^. This expansion may reflect a compensatory epithelial response to impaired AMP production. Tuft cells have emerged as multifunctional chemosensory cells that respond not only to helminths and protists^5–7^, but also to bacterial metabolites^63^. The increased expression of *Sucnr1* and *Vmn2r26* that we observed in tuft cells of TLR2-deficient mice is consistent with an adaptive response that preserves microbial surveillance when ISC-mediated antimicrobial programs are compromised. Whether such responses functionally compensate for impaired Paneth- and enterocyte-derived defense programs under infectious or inflammatory challenge remains to be determined.

These findings may also have implications for intestinal disease. Impaired AMP production is a hallmark of ileal Crohn’s disease, where reduced α-defensin expression has been linked to barrier dysfunction and microbial imbalance^66,67,69^. In addition, genetic and functional studies have associated TLR2 dysfunction with IBD susceptibility^86–88^. Our work suggests that disruption of TLR2 signaling may compromise epithelial defense not only by affecting mature epithelial cells but also by perturbing microbial sensing within the stem cell compartment. This raises the possibility that defective ISC-intrinsic sensing contributes to failure in epithelial regeneration and antimicrobial defense. Whether this pathway is altered in human IBD, and whether restoring TLR2 signaling can repair epithelial function, remain important questions for future study.

Although most of our work was performed in murine systems, our human organoid data indicates that this mechanism is conserved. Human ileal ISC and TA populations express TLR1 and TLR2^75^, and TLR2 stimulation in human ileal primary cultures induced inflammatory target genes and promoted the accumulation of LYZ1^+^ Paneth-like cells. This is particularly relevant given that mature Paneth cells are typically underrepresented in human SI organoid cultures^50,73,74^. TLR2 agonist-based conditioning may therefore provide a useful strategy to improve human organoid models of epithelial differentiation and to study host-pathogen interactions in a more physiologically relevant context. More broadly, the conservation of this pathway supports the translational relevance of ISC-intrinsic microbial sensing.

Taken together, our study identifies ISC-intrinsic TLR2 signaling as a mechanism by which ISCs sense luminal microbial cues and translate them into epithelial differentiation and antimicrobial defense programs. We propose that Lgr5^+^ ISCs maintain host-microbiota symbiosis not only through epithelial renewal, but also by actively detecting microbial signals and adjusting lineage output accordingly. By supporting a role of adult stem cells as microbial sensors, our findings reveal a broader concept in which barrier-tissue stem cells integrate environmental information to coordinate regeneration and host protection. This work opens new avenues for understanding epithelial adaptation and suggests that targeting stem-cell microbial-sensing pathways may provide therapeutic opportunities to restore epithelial homeostasis in intestinal disease.

## Limitations of the study

This study has several limitations. First, although we demonstrate that TLR2 mediates microbial sensing in ISCs, TLR2 is known to recognize a broad range of ligands, including those derived from fungi and viruses. Thus, its role may extend beyond bacterial detection to more generally alert ISCs to changes in the luminal environment, such as intestinal dysbiosis. Second, while TLR2 loss leads to the accumulation of goblet-Paneth intermediate cells, their functional role remains unclear. Third, the expansion of tuft cells, with increased expression of bacterial-sensing receptors, suggests a compensatory mechanism, but their functional contribution to epithelial defense remains to be demonstrated, and further studies are needed to elucidate their role. Fourth, although TLR2-dependent ISC sensing regulates epithelial antimicrobial programs, its impact on microbiome composition was not assessed. Finally, while our data support an epithelial-intrinsic role for TLR2 in ISCs, TLR2 is also expressed by immune cells in the lamina propria, and the role of immune-TLR2 activation was not addressed.

## Acknowledgments

We thank Merav Kedmi for help in RNA single cell experiments and the histology unit. M.B. holds the Ernst and Kaethe Ascher Career Development Chair. This study was supported by research grants from the Center for New Scientists at the Weizmann Institute of Science, the Israel Science Foundation (grant No. 1587/20 and 3775/25), the Helen and Martin Kimmel Institute for Stem Cell Research at The Weizmann Institute of Science, and the Minerva Foundation, with funding from the Federal German Ministry for Education and Research, the Moross Integrated Cancer Center, the Israel Ministry of Science (IMOS, Grant No. 4631), the Dr. Gilbert S. Omenn and Martha A. Darling Weizmann Institute - Schneider Hospital Fund for Clinical Breakthroughs through Scientific Collaborations, a research grant from the Snider Foundation, the Abisch-Frenkel RNA Therapeutics Center, a research grant from the Shimon and Golde Picker, and a research grant from the Herbert K. Bennett Charitable Fund, Dwek Institute for Cancer Therapy Research.

## Author contributions

A.H.M. and M.B. conceived the study, designed experiments, and interpreted the results; C.K. and A.S.P. designed and performed the computational analysis with the assistance of B.T. and L.A.; A.H.M. carried out all experiments with the help of S.L., V.H., C.S., and N.D.; S.L. performed RNA single-cell experiments; V.H. assisted with GF colonization experiments; S.L.Z., N.D. and L.O performed the TEM experiments; D.D. and M.G. provided and assisted with human organoid experiments; I.K. provided human biopsies used for organoid production; M.B. supervised this study; A.H.M., C.K., L.A., and M.B. wrote the manuscript, with input from all authors.

## Methods

### Mice

All mouse work was performed in accordance with the Institutional Animal Care and Use Committees (IACUC) and relevant guidelines at the Weizmann Institute of Science (IACUC, no. 04270724-1). C57BL/6J wild type (WT), TLR2 cKO (strain name: B6.129-Tlr2tm1Kir/J; stock number 004650), TLR4 cKO (strain name: B6(Cg)-Tlr4tm1.2Karp/J; stock number: 029015), CD45.1 (strain name: B6.SJL-Ptprca Pepcb/BoyJ; stock number: 002014), GFP (strain name: C57BL/6-Tg(UBC-GFP)30Scha/J; stock number: 004353) and Villin-Cre (strain name: B6.Cg-Tg(Vil1-cre)997Gum/J; stock number: 004586) mice were purchased from the Jackson Laboratory. Lgr5-2A-eGFP reporter mice were gifted by Barker N. group (A*Star Institute of Medical Biology) and were described before^45^. Lgr5-GFP labeled TLR2 cKO mice were generated by crossing TLR2 cKO with Lgr5-2A-EGFP. TLR2^fl/fl^ mice were a generous gift from Prof. Clifford V. Harding (Case Western Reserve University). TLR2^ΔIEC^ mice were generated by crossing TLR2^fl/fl^ with Villin-Cre. TLR2^ΔISC+tdTomato^ mice were generated by crossing TLR2^loxp^ (TLR2^fl/fl^) with Lgr5-2A-CreERT2; Rosa26 LSL-tdTomato mice. Germ-free (GF) C57BL/6 mice were born in the Weizmann Institute germ-free facility and routinely monitored for sterility. Both male and female age-matched mice, ranging from 8 to 14 weeks of age, were used for all experiments in this study. Littermates of the same genotype, sex, and age were randomly assigned to experimental groups. All mice, except GF, were housed under specific-pathogen-free (SPF) conditions at the Weizmann Institute animal facilities.

### Generation of BM-chimeric reconstituted mice

To generate chimeric mice, 8-week-old TLR2-sufficient Lgr5-GFP mice (chimeric control) and TLR2 cKO crossed with Lgr5-GFP mice (chimeric KO) were irradiated twice with 500 rad per mouse at an interval of 2h (X-RAD 320 X-Ray Irradiator). Mice were kept for two weeks on broad-spectrum antibiotics in their drinking water (Enrolfoxacin 1:200). One day later, BM mononuclear cells were isolated from donor (GFP or CD45.1) mice, as indicated in each experiment, by flushing the femur bones. Red blood cells were lysed with ACK lysing buffer (Gibco). BM cells were resuspended in PBSx1 and a total of 5 million cells per mouse. BM cells were injected retro-orbitally into the irradiated mice. After 4 weeks, peripheral blood samples were collected and analyzed by FACS to assess reconstitution in the blood or lamina propria.

### Lineage tracing

To label Lgr5+ ISC and their following progeny for *in vivo* lineage tracing, we administered one intraperitoneal injection of Tamoxifen (2 mg per 20 g body weight) to induce Cre-mediated excision of a stop codon and subsequent expression of tdTomato from the Rosa26 locus.

### Epithelial cell dissociation and crypt isolation

For all mice, crypts were isolated from the distal part of the SI. The SI was extracted and rinsed in cold phosphate-buffered saline (PBS). The tissue was opened longitudinally and sliced into small fragments, approximately 0.2 cm long, followed by incubation with 20 mM EDTA-PBS on ice for 90 minutes. Then, the tissue was shaken vigorously, and the supernatant was collected as fraction 1 in a new conical tube. The tissue was incubated in PBS, and a new fraction was collected every 5 min. Fractions were collected until the supernatant consisted almost entirely of crypts. The fractions 3-4 (enriched for crypts) were filtered through a 70µm filter, washed twice in PBS, centrifuged at 400g for 5 min, and dissociated with TrypLE Express (Gibco) for 1 min and 30 seconds at 37°C. The single-cell suspension was then passed through a 40μm filter and stained with fluorescence-activated cell sorting (FACS) antibodies and sorted with SH800 Sony sorter for subsequent analysis (bulk or single cell RNA-sequencing and qPCR).

### Flow cytometry analysis

Single-cell suspensions were prepared as described above. Epithelial cells were stained EpCAM (Biolegend, G8.8), CD45 (Biolegend, 30-F11), CD24 (Biolegend, M1/69), human Lyz1-FITC (Dako, EC 3.2.1.17), and Dapi-NucBlue (ThermoFisher, cat no. R37605) for 30 minutes on ice. Analysis was done on a Cytoflex analyzer or a Sony sorter. Further analysis of the data was done using FlowJo v10.1.

### Cell sorting

For the bulk population, a sorter FACS (SH800 Sony) was used to sort 1000-10,000 cells into an Eppendorf tube containing 50μl of TCL buffer (QIAGEN) solution with 1% 2-mercaptoethanol (Sigma Aldrich). To enrich for Lgr5^+^ ISC population, cells were isolated from Lgr5-2A-eGFP mice, stained with the antibodies CD45 (Biolegend, 30-F11), EpCAM (Biolegend, G8.8), CD24 (Biolegend, M1/69), Lyz1 (Dako, EC 3.2.1.17) and Dapi-NucBlue (ThermoFisher, cat no. R37605). Cells were gated for EpCAM^+^ CD45^-^ (IEC), EpCAM^+^ CD45^-^ GFP^high^ (Lgr5+ ISC) or EpCAM^+^ CD45^-^ CD24^+^ FSC^high^ (Paneth cells). Lineage-traced cells were identified based on endogenous tdTomato fluorescence. Cells were gated for EpCAM^+^ CD45^-^ tdTomato^+^. For bulk RNA-sequencing, the tubes were centrifuged and immediately frozen on dry ice and kept at 80°C until ready for RNA isolation. For scRNA-sequencing, the cells were treated as described below.

### Stimulation of flow-sorted intestinal stem cells

Intestinal epithelial single-cell suspensions were prepared from distal SI as described before. ISCs were sorted on a Sony SH800 (Lgr5–eGFP^high^). Immediately after sorting, cells were pelleted (300 ×g, 5min at 4°C) and resuspended in 100µl L-WRN C.M. supplemented with 10µM Y-27632. TLR2 agonist Pam3Csk4 (1μg/mL, InvivoGen) was added and incubated for 4h at 37 °C, 5 % CO₂, static. Cells were pelleted and resuspended in TCL buffer containing 1% 2-mercaptoethanol. The RNA was then cleaned and amplified using the Smart-Seq2 library preparation protocol.

### Hash-tagged single-cell RNA-sequencing

Single-cell RNA-sequencing (scRNA-seq) libraries were prepared using the Chromium single-cell RNA-seq platform (10x Genomics). Isolated IEC were sorted into a cooled 15 mL tube with 0.04% BSA in PBS using a Sony SH800 cell sorter. Cells for each condition or mouse were stained using Biotin anti-mouse CD326 (EpCAM) antibody (cat #118204) followed by staining using TotalSeq™ PE Streptavidin (B0952 - B0955). 20,000 cells were sorted for each mouse or condition and pooled following the manufacturer’s instructions. 30,000 single-cell suspension was loaded onto Next GEM Chip G targeting 15,000 cells and then ran on a Chromium Controller instrument to generate GEM emulsion. Single-cell 3’ RNA-seq libraries and cell surface protein libraries were generated according to the manufacturer’s protocol (Chromium Single Cell 3’ Reagent Kits User Guide (v3.1 Chemistry Dual Index)). Final libraries were quantified using NEBNext Library Quant Kit for Illumina (NEB) and high-sensitivity D5000/D1000 TapeStation (Agilent). Libraries were pooled according to targeted cell number, aiming for ∼20,000 reads per cell for gene expression libraries and ∼5,000 reads per cell for cell surface protein libraries. Pooled libraries were sequenced on a NovaSeq 6000 instrument using an S1 100 cycles reagent kit (Illumina).

### Bulk population RNA purification and cDNA preparation

Libraries were prepared using a modified SMART-Seq2 protocol as previously reported^89^. RNA lysate cleanup was performed using RNAClean XP beads (Agencourt), followed by reverse transcription with Maxima Reverse Transcriptase (Life Technologies) and whole transcription amplification (WTA) with KAPA HotStart HIFI 2 3 ReadyMix (Kapa Biosystems) for 18 cycles. WTA products were purified with Ampure XP beads (Beckman Coulter), quantified with Qubit dsDNA HS Assay Kit (ThermoFisher), diluted to a concentration of 1 ng/μL in ultra-pure water, and assessed with a high-sensitivity DNA chip (Agilent). RNA-seq libraries were constructed from purified WTA products using the Nextera XT DNA Library Preparation Kit (Illumina). The libraries were sequenced on an Illumina NextSeq 500/550 high kit.

### Quantitative RT-PCR

For quantitative PCR (qPCR), cDNA from bulk population libraries were diluted to a final concentration of 0.01ng/μl. Fast SYBR green master mix (Thermo-Fisher scientific) was used to perform qRT–PCR per the manufacturer’s instructions. The primer sequences are provided in **Supplementary Table 10**. Data analysis was performed with QuantStudio 12K flex software (Thermo-Fisher scientific) based on the ΔΔCT method. All target genes were standardized to the reference gene Ubiquitin C (UBC).

### Murine intestinal stem-enriched spheroid cultures

Small intestinal crypts were isolated from 8–12-week-old wild-type C57BL/6J or TLR2 cKO mice. After two PBS washes (300g for 5 min at 4 °C), the crypt number was estimated by bright-field microscopy (EVOS™ M5000 Imaging System). Crypts were pelleted, and roughly 500 crypts were embedded with 20µl ice-cold Matrigel™ (Corning) containing 1 µM Jagged-1 peptide (AnaSpec, AS-65155) and plated as a dome in a pre-warmed 24-well plate. After polymerisation (20 min, 37°C), spheroid domes were overlaid with 600µl of 50% L-WRN ^32^ conditioned medium supplemented with 10µM Y-27632 (Biogems, 1293823). Fresh L-WRN conditioned medium was replaced one day after seeding, and spheroids were passaged by dissociation with 0.05% Trypsin/0.5 mM EDTA (Bio-Lab Ltd) and resuspended in new Matrigel on day 3 with a 1:2 split ratio. Alternatively, for differentiation, organoid domes were overlaid with 600µl organoid culture medium (Advanced DMEM/F12, Invitrogen) with streptomycin/penicillin and GlutaMAX and supplemented with EGF (150 ng/mL, Peprotech), R-Spondin-1 (625ng/mL, Peprotech), Noggin (100ng/mL, Peprotech), Y-276432 (10μM, biogems), N-acetyl-1-cysteine (1μM, Sigma-Aldrich), N2 (1X, Gibco), B27 (1X, Gibco). Fresh media was replaced on day 3, and organoids were passaged by mechanical dissociation and re-suspended in new Matrigel on day 6 with a 1:3 split ratio.

For TLR2 stimulation, the TLR2 agonist Pam3Csk4 (1μg/mL, InvivoGen) was added to the growth medium 4 and 8 hours prior to RNA harvesting. Each well was recovered separately from Matrigel by washing with PBS. The sample pellet was resuspended in TCL buffer containing 1% 2-mercaptoethanol, and then the RNA was cleaned and amplified using the Smart-Seq2 library preparation protocol.

### Formation of transwell stem-enriched monolayers

To form stem-enriched epithelial monolayers ^39^, spheroids were taken from 3-day-old 3D cultures for plating in Transwells (3470; Corning Costar). The Transwells were coated in 10% Matrigel™ for 30 minutes at 37 °C. Spheroids were recovered from Matrigel by first washing in PBS and then dissociated for 3 min at 37 °C using a solution of 0.05% Trypsin/0.5 mM EDTA (Bio-Lab Ltd). The trypsin was then inactivated using DMEM media (Gibco) containing 10% fetal bovine serum. The spheroids were then dissociated by vigorous pipetting (using a 1,000 μl pipette). The cells were then passed through a 40 μm cell strainer and re-suspended in 50% L-WRN containing 10 μM Y-27632 (Biogems, 1293823). Cells were plated into the upper compartment of a 6.5 mm Transwell with 0.4 µm Pore Polyester Membrane Insert (Corning) at approximately 3-4x10^4^ cells/transwell in 200 μl of media. An additional 800 μl of media was added to the lower compartment of the Transwells. The medium was replaced with 50% L-WRN conditioned medium containing 10µM Y-27632 in both the top and bottom compartments and replenished every 2 days.

For TLR2 stimulation, the TLR2 agonist Pam3Csk4 (1μg/mL, InvivoGen) was added to upper or lower growth medium 4 and 24 hours prior to RNA harvesting. Each well was recovered separately from Matrigel by washing with PBS. The sample pellet was resuspended in TCL buffer containing 1% 2-mercaptoethanol, and then the RNA was cleaned and amplified using the Smart-Seq2 library preparation protocol.

### Polarization of stem-enriched monolayers on transwell inserts

To demonstrate that the seeded cells formed a functional monolayer, we measured transepithelial electrical resistance (TEER) in the Transwells every 3 days. The average TEER of untreated cells was 500-600 Ωcm^2^. The TEER was tested as indicated with an EVOM3 apparatus (World Precision Instruments).

### FITC-Dextran-uptake assay

The diffusion was measured by the addition of FITC-labelled dextran (Sigma-Aldrich; 4 kDa) to the apical surface at a final concentration of 2 mg /ml. After 2h incubation at 37 °C, 100 µl aliquots of the basal media were collected, and the fluorescence was measured at 495 nm using Cytation 5 plate reader (BioTek). As a positive control, the fluorescence of a cell-free Matrigel-coated transwell was measured to assess the diffusion of FITC-labelled dextran.

### Human organoid assay

Human ileal organoids used in this study were obtained from the Israeli Hadassah Organoid Center (HOC) and generated from biopsies collected from the distal ileum. The collection and use of these organoids were approved under the Committee on Research Involving Human Subjects application (0921-20-HMO, The Israeli Organoid Bank- A Means For- Personalised Medicine) and the Weizmann Institute of Science IRB.

In brief, organoids were cultured in IntestiCult Organoid Growth Medium (Human, Stemcell, Cat no. 06010) in 3D Matrigel in 24-well plates. In experiments requiring differentiation, IntestiCult Differentiation Medium (Human, Stemcell, Cat. No. 100-0214) was used. Organoids were stimulated at different time points with TLR2 agonist Pam3Csk4 (1μg/mL, InvivoGen) in IntestiCult Organoid Growth Medium/differentiation Medium before RNA extraction or imaging. Organoids were stimulated, washed, fixed, and stained in an Ibidi chamber for imaging.

### Histochemistry

Tissues were fixed for 16-24 hours in formalin, embedded in paraffin, and cut into 5μm thick sections. Sections were deparaffinized with standard techniques. For H&E staining, slides were stained with Hematoxylin for 1 minute, washed, stained with Eosin for 45 seconds. For PAS and Alcian blue staining, tissue was stained with Alcian blue and periodic acid-Schiff reagents kit (ScyTek Laboratories, cat no. APS-1) on deparaffinized slides with standard techniques, using 3% acetic acid for 3 minutes, then Alcian blue solution (Alcian blue, pH 2.5) for 30 min, rinsed in tap water for 5 min, and oxidized in periodic acid (0.5%) for 10 min, followed by rinsed in running tap water for 5 min, and stained in Schiff reagent as a counterstain (cancer diagnostics) for 10 min. Nuclei were stained with PureView™ Mayers Hematoxylin for 50 seconds. All slides were washed, dehydrated and mounted with Sub-x mounting medium (Leica).

### Immunofluorescence (IF)

Tissues were fixed for 14 hours in formalin, embedded in paraffin, and cut into 5 μm thick sections. Sections were deparaffinized with standard techniques, incubated with primary antibodies overnight at 4°C (**Supplementary Table 10**), followed by secondary antibodies incubation (**Supplementary Table 10**) at room temperature for 30 min. Slides were mounted with Slowfade Mountant+DAPI (Life Technologies, S36964) and sealed.

### Single-molecule fluorescence in situ hybridization (smFISH)

RNAScope Fluorescent Multiplex V2 kit (Advanced Cell Diagnostics) was used according to the manufacturer’s recommendations with the following alterations. The target retrieval boiling time was adjusted to 16 minutes, and the incubation with Protease IV at 40°C was adjusted to 7 minutes. Slides were mounted with Slowfade Mountant+ DAPI (Life Technologies, S36964) and sealed. Probes used for single-molecule RNAscope (Advanced Cell Diagnostics) are described in **Supplementary Table 10.**

### Combined immunofluorescence and smFISH

Following smFISH protocol as described above, tissue sections were washed in washing buffer, incubated with primary antibodies overnight at 4°C, washed in 1x TBST 3 times, and then incubated with secondary antibodies for 30 min at room temperature. Slides were mounted with Slowfade Mountant+DAPI (Life Technologies, S36964) and sealed.

### TUNEL assay

Small intestine tissue was analyzed for TUNEL staining by the TUNEL Assay Kit (Abcam cat no. 1187 ab66110) according to the manufacturer’s recommendations. In brief, sections were deparaffinized followed by antigen retrieval, labeled with BrdU for 1 hour at 37°C, and incubated with anti–BrdU-Red antibody for 30 minutes at room temperature. After TUNEL labeling, the sections were mounted with hoechst Blue (1 µg/mL) (Invitrogen R37605) for 5 minutes at room temperature, followed by a PBS wash. (. Measurement and analysis of the staining area was performed using Zen software (Zeiss).

### Image analysis

Images of tissue sections were taken with a confocal microscope, LSM900 (Zeiss). Scale bars were added to each image using the confocal Zen analysis software (Zeiss). Images were overlaid and visualized using ImageJ software and Zen analysis software (Zeiss).

### Transmission electron microscopy

Tissues were fixed with 4% paraformaldehyde, 2% glutaraldehyde in 0.1 M cacodylate buffer containing 5 mM CaCl_2_ (pH 7.4), then postfixed with 1% osmium tetroxide supplemented with 0.5% potassium hexacyanoferrate tryhidrate and potasssium dichromate in 0.1 M cacodylate (1 hour), stained with 2% uranyl acetate in water (1 hour), dehydrated in graded ethanol solutions and embedded in Agar 100 epoxy resin (Agar scientific Ltd., Stansted, UK).

Ultrathin sections (70-90 nm) were viewed and photographed with a FEI Tecnai SPIRIT (FEI, Eidhoven, Netherlands) transmission electron microscope operated at 120 kV and equipped with a OneView Gatan Camera.

### Image quantification

Quantification of IFA and smFISH images from all tissues was assessed by staining for E-cadherin (BD Biosciences) to mark cell borders and DAPI staining for nucleus visualization. Cells were manually counted based on immunofluorescence staining specific to each cell type. Mature Paneth cells were similarly quantified using Lyz1 (Dako) and Mptx2 (prob (ACDBio) or antibody (Abcam)) as markers. Tuft cells were quantified along the villi-crypt axis using DCLK1 (Abcam) marker. Tuft2 cells were also quantified using the CD45 (Biolegend) marker. Proliferative cells (Ki67+) were counted along the villus-crypt axis. For each quantification, at least ten intact and longitudinally oriented crypts per tissue section were analyzed.

Quantification of histochemistry images of goblet-paneth intermediate cells was assessed by AB-PAS staining in different models within the crypts. Mature Paneth cells were quantified based on their presence of PAS staining. Goblet-Paneth intermediate cells were assessed by AB-PAS staining. For each quantification, at least ten randomly intact and longitudinally oriented crypts per tissue section were analyzed.

### Microbiota isolation and colonization Microbiota isolation

Luminal bacteria isolation was performed under anaerobic conditions using an anaerobic chamber. Fresh intestinal contents were collected from adult mice (C57BL/6J WT, and TLR2 cKO, *n*=2 each). Mice were euthanized, and the SI and cecum were immediately harvested inside the chamber. Luminal contents were gently collected by flushing without directly handling or compressing the intestinal tissue. Large food particles and debris were removed by centrifugation at 300 × g for 5 minutes. The supernatant was centrifuged at 4,000 × g for 15 minutes to pellet the bacterial fraction. The bacterial pellets were resuspended in 1 mL of sterile PBS containing 30% glycerol.

### Colonization protocol

Bacterial colonization was performed in a gnotobiotic-like workflow using sterile isocages. Two experimental groups (*n*=4 mice/group) were housed in individual isocages prior to colonization. Food and water were withdrawn 3 hours before colonization. All colonization procedures were conducted in a biosafety cabinet. Sterile, autoclaved oral gavage needles were used (one per group). The gavage needle was inserted along the left side of the oral cavity at a slight angle to minimize trauma. Each mouse received 100μL of freshly mixed microbiota suspension via oral gavage. Successful delivery was confirmed by the absence of reflux after needle removal.

Following colonization, water was returned to the cages immediately, and food was restored an hour later. The colonization procedure was repeated on days 3 and 6 post-initial colonization using 30% glycerol-frozen bacterial pellets from the initial microbiota isolation. Mice were harvested on day 14 post-colonization for analysis.

### Statistical analysis

Statistical analysis was performed using Prism (GraphPad). R v4.4.1, R v4.3.1, and R2020a. The specific tests applied are detailed in the corresponding Fig. legends and computational methods.

### Computational analysis

#### Bulk RNA sequencing

Poly-A/T stretches and Illumina adapters were trimmed from the reads using cutadapt^90^; resulting reads shorter than 30bp were discarded. Reads were mapped to the *M. musculus* reference genome GRCm39 using STAR^91^, supplied with gene annotations downloaded from Ensembl (and with EndToEnd option and outFilterMismatchNoverLmax was set to 0.04). Expression levels for each gene were quantified using htseq-count ^91^, using the gtf above. Differentially expressed genes were identified using DESeq2 ^92^ with the betaPrior, cooksCutoff and independent Filtering parameters set to False. Raw P values were adjusted for multiple testing using the procedure of Benjamini and Hochberg. Pipeline was run using snakemake^93^. Differentially expressed genes were determined by a p-adj of < 0.05 and absolute fold changes > 2. Heatmap plotting was done using the ComplexHeatmap package from R (Gu Z (2015). *ComplexHeatmap: Making Complex Heatmaps*. R package version 1.0.0, https://github.com/jokergoo/ComplexHeatmap.) Go enrichment was done using ClusterProfiler package in R^94^ for up and down genes separately.

Published dataset of smart-seq2 sequencing^4^ (accession number GSE92332) was used to acquire the expression of TLRs on IECs.

#### Single-cell RNA-Seq processing

Cell Ranger (version 7.1.0) was used for demultiplexing and alignment to the reference genome GRCm38 (mm10). Consequently, 11,864 (70.57%) cells could be assigned to a single sample with high confidence. Cells considered as low quality were removed when one of the following criteria was met: expression < 1500 genes, percentage of mitochondrial gene counts above 40% or predicted doublet by scrublet. 6243 cells passed the filtering steps. Downstream analysis was performed using Python version 3.9.18 and the methods of scanpy (version 1.10.13) and scvi-tools (version 1.1.6.post2) with default parameters if not mentioned otherwise. First, 3000 highly variable genes were selected using the Seurat v3 approach. Next, a reduced, latent representation (30 dimensions) of each cell was learned by the single-cell Variational Inference (scVI) model of scvi-tools ^95^. Samples were provided as a batch for correction and the percentage of mitochondrial gene counts and the library size as covariate variables. The model was trained on a batch size of 127 and early stopping was triggered after 113 epochs. The latent space was used to calculate a neighborhood graph with subsequent cluster identification using leiden algorithm with a resolution of 0.8. Uniform manifold approximation and projection (UMAP) and diffusion maps were applied to gain a low-dimensional representation of each cell for visualization purposes. The differential expression analysis was based on the Wilcoxon rank sum test.

To characterize mature enterocytes as having either a proximal or distal phenotype, we utilized the published scRNA-seq dataset from Haber et al.^4^ as a reference. The Pearson correlation coefficient (scipy version 1.11.4) was calculated between the raw counts of each cell within the mature enterocyte cluster and the median raw values of the “Mature Distal Enterocytes” and “Mature Proximal Enterocytes” clusters from the reference dataset, respectively.

To quantify the spatial identity of each cell along the crypt-villus axis, we calculated the zonation score based on the landmark genes by Moor et al.^53^. This score was determined for each cell by summing the normalized expression of landmark genes for the intestinal villus tip and, separately, for the intestinal crypts. The final zonation score was computed as the ratio of the aggregated villus tip expression to the sum of both aggregated villus tip and crypt expression values.

### Screening for encrypted antimicrobial peptides

To identify potential encrypted antimicrobial domains, a reference database of known validated and predicted AMPs was constructed, using amino acid sequences from public databases (CAMPR4, DRAMP, ribosomal and non-ribosomal DBAASP, dbAMP). The integrated AMP reference database was further filtered to exclude any unspecific sequences shorter than 5 or 12 amino acids (**Supplementary Table 8**). The protein sequences of murine target genes were retrieved via the UniProt REST API. The resulting protein sequences were downloaded in FASTA format and parsed into a structured dataframe using the Biopython library. Encrypted AMPs within the target proteins were identified by an exact substring match with sequences of the reference database.

For every matched AMP sequence an absolute “Torres” score was calculated^56^. Biophysical properties (i.e. net charge, hydrophobicity) were calculated using the Bio.SeqUtils.ProtParam.ProteinAnalysis module from the Biopython (version 1.86) library.

## Supplementary Fig. legends

**Extended Data Fig.1.**
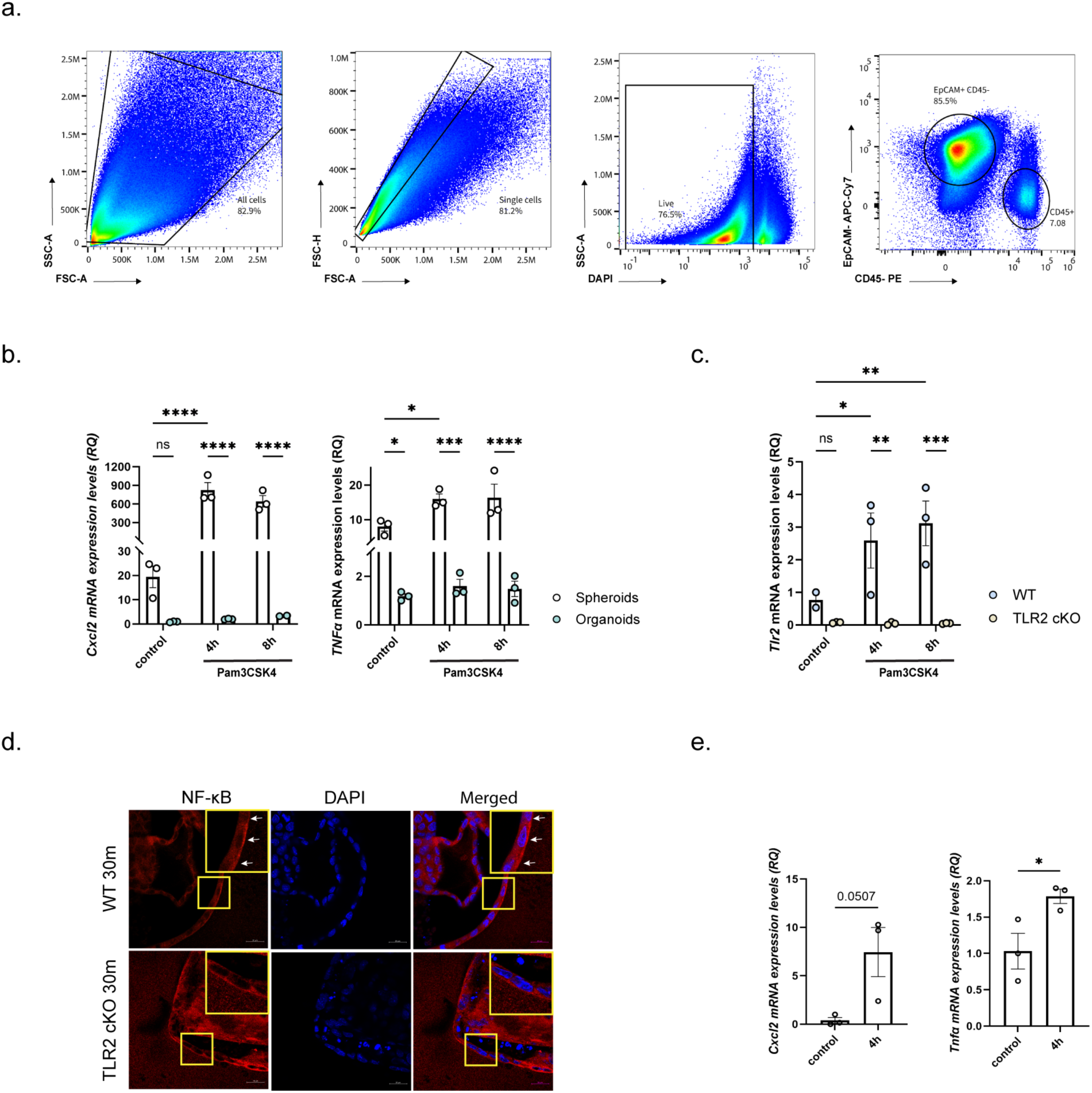
The TLR2 pathway is functional in stem cells. (**a**) A representative flow cytometry gating strategy for epithelial cells. (**b**) Relative quantification (RQ) of *Cxcl2* and *Tnfα* expression levels in spheroids vs. organoids generated from WT mice after 4h or 8h of *ex vivo* activation with TLR2/TLR1 agonist (Pam3Csk4, 1μg/mL) or without (control) using qPCR. Data are presented as mean ± SEM. *n*=3 biological repetitions per group, \**p* <0.05, \*\*\**p* <0.001, \*\*\*\**p*< 0.0001, two-way ANOVA. (**c**) Relative quantification (RQ) of *Tlr2* expression in spheroids generated from WT or TLR2 cKO mice after 4h or 8h of *ex vivo* activation with TLR2/TLR1 agonist (Pam3Csk4, 1μg/mL) or without (control) using qPCR. Data are presented as mean ± SEM. *n*=3 biological repetitions per group, except for the control time 0 with 2 biological repetitions, \**p* <0.05, \*\**p* <0.01, \*\*\**p* <0.001, two-way ANOVA. (**d**) NFĸB pathway activation. Representative images of IFA of NFĸB subunit (p65, red) and DAPI in WT (upper panel) and TLR2 cKO (lower panel) stem-enriched spheroids after 30 minutes of Pam3Csk4 (1μg/mL) induction. Scale bar, 20μm. Insets, x2 magnification. (**e**) Relative quantification (RQ) of *Cxcl2* and *Tnfα* expression levels in isolated Lgr5^+^ ileal stem cells from WT mice after 4h of *ex vivo* activation with TLR2/TLR1 agonist (Pam3Csk4, 1μg/mL) or without (control) using qPCR. Data are presented as mean ± SEM. *n*=3 biological repetitions, \**p* <0.05, two-tailed *Student’s t-test*.

**Extended Data Fig.2.**
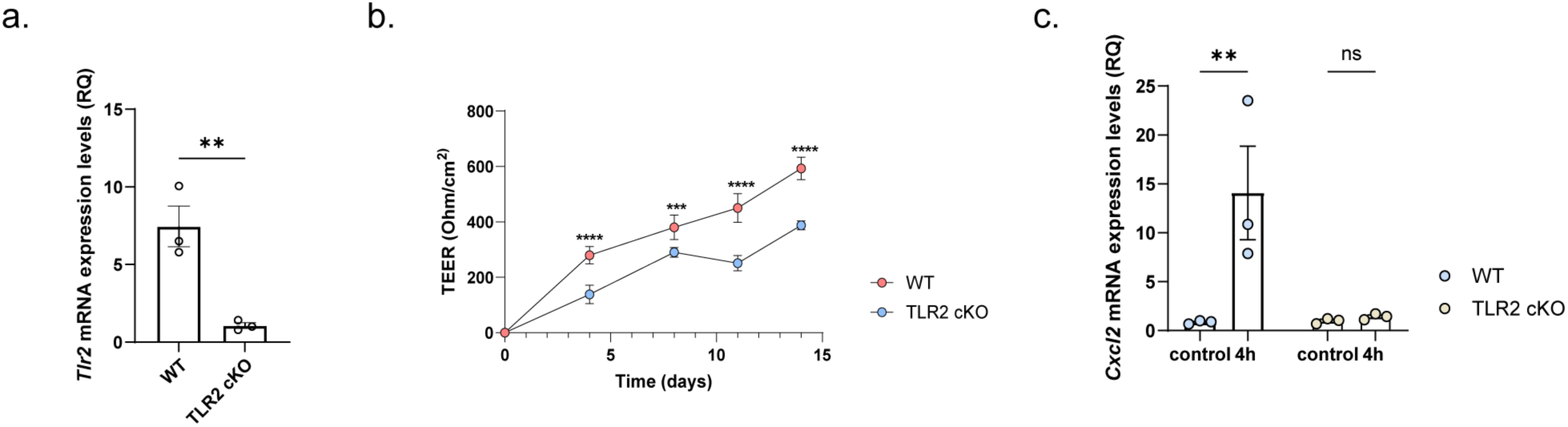
Polarized TLR2 activation in stem-enriched monolayers. (**a**) Relative quantification (RQ) of *Tlr2* expression in stem-enriched monolayers derived from WT vs TLR2 cKO using qPCR. Data are presented as mean ± SEM. *n*=3 biological repetitions, \*\**p* <0.01, two-tailed *Student’s t-test*. (**b**) Monolayer barrier integrity assay using TEER measurements of WT or TLR2 cKO. Data are presented as mean ± SEM. *n*=3 biological repetitions; \*\*\**p* <0.001, \*\*\*\**p*< 0.0001, two-tailed *Student’s t-test*. (**c**) Relative quantification (RQ) of *Cxcl2* expression levels in stem-enriched monolayers from WT or TLR2 cKO mice after 4h of activation with TLR2/TLR1 agonist (Pam3Csk4, 1μg/mL) or without (control) using qPCR. Data are presented as mean ± SEM; *n*=3 biological repetitions; *ns*, \*\**p*<0.01, two-way ANOVA.

**Extended Data Fig. 3.**
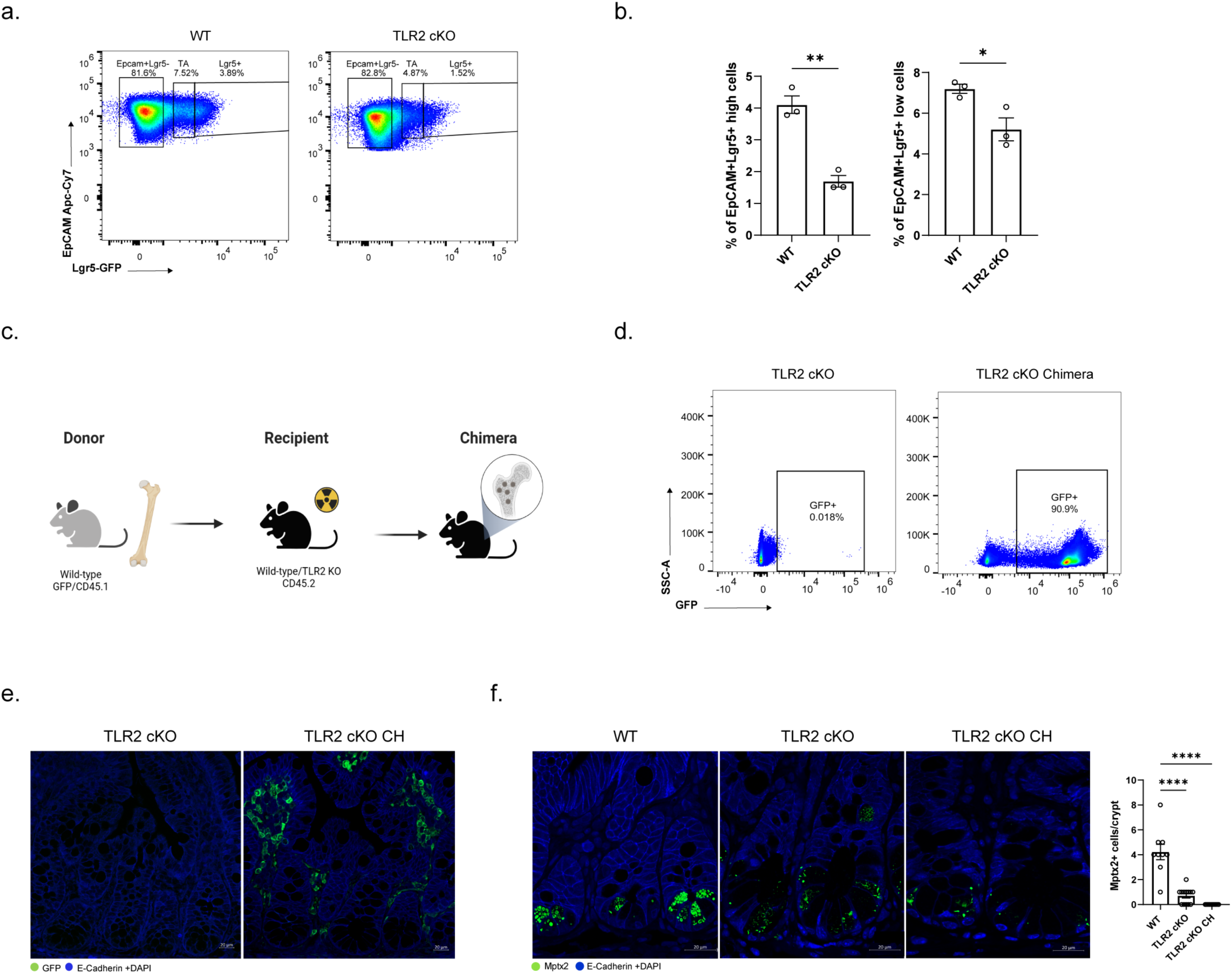
TLR2 KO chimeric mice model characterization. (**a-b**) Representative flow cytometry plot (a), and quantification of the percentage of Lgr5^+^ ISC GFP^high^ (left) and GFP^low^ (right) EpCAM^+^ cells from ileal crypts in WT or TLR2 cKO (b). Data are presented as mean ± SEM. *n*=3 mice per group; \**p*< 0.05, \*\**p*< 0.01, two-tailed *Student’s t-test.* (**c**) schematic representation of the TLR2 KO chimeric mice model. TLR2 cKO mice were irradiated with a 500 Rad dose twice, followed by bone marrow transplantation (BMT) from GFP/CD45.1 mice 24h later. (**d-e**) Confirmation of BMT in the TLR2 cKO chimeric mouse model. (d) Representative flow cytometry plot of GFP-labeled immune cells in the LP. (e) immunofluorescence assay (IFA) of GFP (green), E-cadherin (blue), and DAPI from TLR2 cKO or chimeric TLR2 cKO (TLR2 cKO CH) mice. E-Cadherin (blue) stains IEC boundaries. Scale bar, 20μm. (**f**) Mptx2 staining in Paneth cells from SI ileal crypts of WT (left), TLR2 cKO (mid), and chimeric TLR2 cKO (TLR2 cKO CH). IFA of Mptx2 (green), E-Cadherin (blue), and DAPI. E-Cadherin (blue) stains IEC boundaries. Scale bar, 20μm. Cells with Mptx2+ granules were quantified per crypt (right); 10 crypts per mouse. Data are presented as mean ± SEM. \*\*\*\**p* < 0.0001, one-way ANOVA.

**Extended Data Fig. 4.**
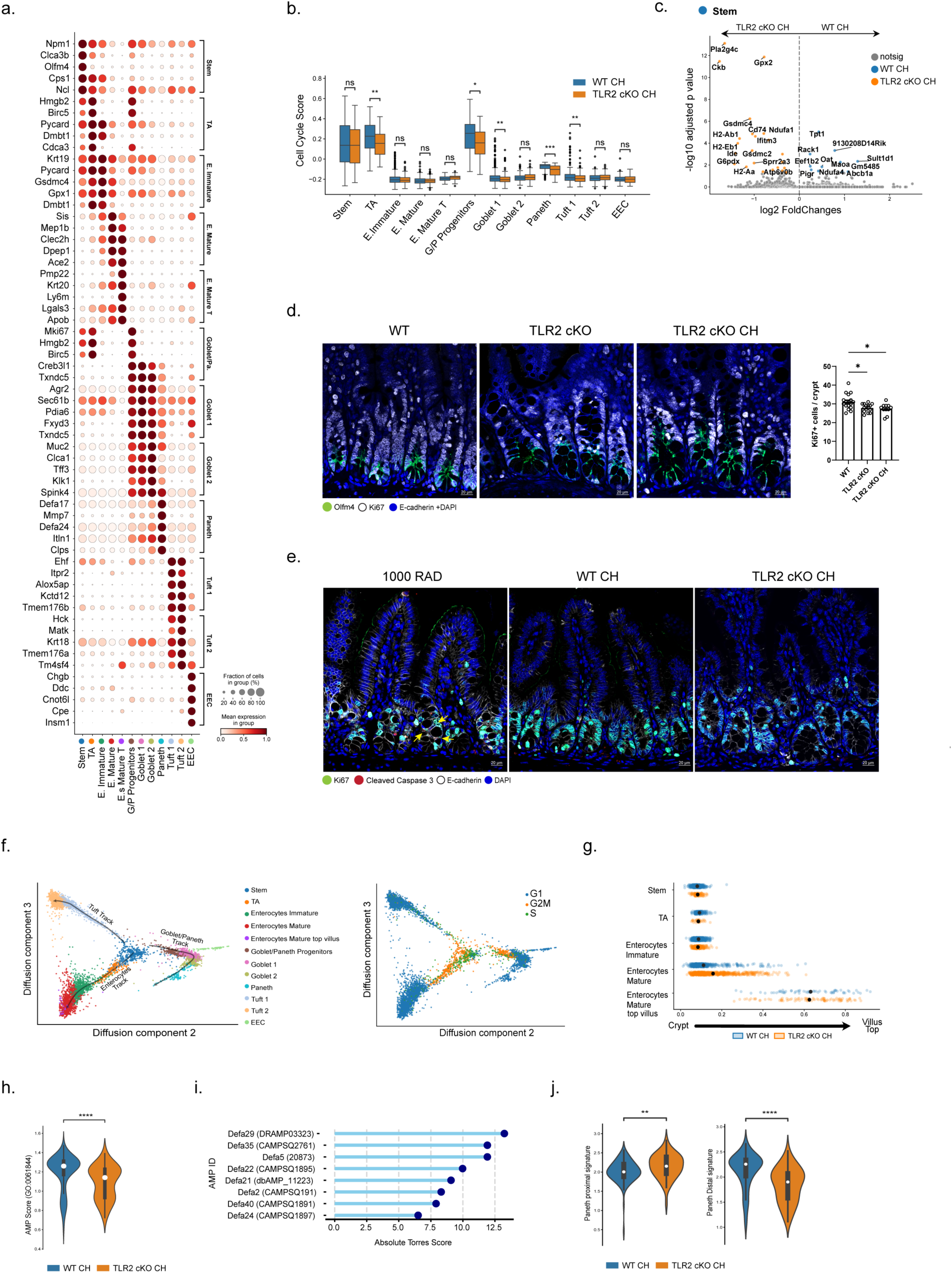
Characterization of TLR2 deletion at single cell resolution. (**a**) Epithelial cell subset markers used for cluster annotation. Dot plot of five known markers per epithelial cell subset derived from publicly available data^4^. The size of the dot indicates the proportion of cells expressing a gene, while the color indicates the mean expression level. TA, transit amplifying; E. Immature, immature enterocyte; E. Mature, mature enterocyte; E. Mature T, mature top-villus enterocytes; G/P Progenitors, goblet-Paneth progenitors; EEC, enteroendocrine. (**b**) Box-and-whisker plot of cell cycle signature score in different epithelial clusters from chimeric (CH) WT (blue) vs TLR2 cKO (orange) mice. *n*=2 per condition. EEC, enteroendocrine. Boxes represent median and Interquartile Range (IQR); whiskers extend to 1.5× IQR; ns (non-significant), \**p* < 0.05, \*\**p*<0.01, \*\*\**p*<0.001, *Wilcox-test*. (**c**) Volcano plot showing log2fc estimates and the negative log10 of enrichment padj. of top upregulated in WT (blue, right) and downregulated (orange, left) DEG in stem cells compared to TLR2 cKO (*n*=2 mice per condition). Colored points correspond to padj ≤ 0.05 and |log2FoldChange| ≥ 1. Color coding is shown in the top-right corner. (**d**) Proliferation analysis in SI ileal crypts of WT (left panel), TLR2 cKO (middle panel), and TLR2 cKO chimera (right panel). IFA of Ki67 (white), Olfm4 (green), E-Cadherin (blue), and DAPI. E-cadherin (blue) stains IEC boundaries. Scale bar, 20μm. Ki67^+^ Cells were quantified per crypt (right); 10 crypts per mouse. Data are presented as mean ± SEM. \**p*<0.05, one-way ANOVA. (**e**) Apoptosis analysis in SI ileal crypts of positive control (24h post-irradiation, left), chimeric (CH) WT (middle), or TLR2 cKO (right) mice. IFA of Cleaved caspase 3 (CC3) (red), Ki67 (green), E-cadherin (white) and DAPI. E-cadherin (white) stains IEC boundaries. Yellow arrows show CC3+ cells. Scale bar, 20μm. (**f**) Diffusion map of differentiation trajectory to the main epithelial lineages colored by cell subset (left) or by cell cycle state (right) in chimeric (CH) WT and TLR2 cKO. Color coding is shown on the right. TA, transit amplifying; EEC, Enteroendocrine. (**g**) Diffusion pseudotime (dpt) score comparing the enterocyte subsets differentiation state of chimeric (CH) WT (blue) and TLR2 cKO (orange). Dpt scores showing an accelerated differentiation in TLR2 cKO CH. (**h**) Distribution of Antimicrobial humoral immune response mediated by antimicrobial peptide-signature score in Paneth cells from chimeric WT vs. TLR2 cKO; significance was determined by Wilcox-test, \*\*\*\**p*<0.0001. (**i**) Top AMP-encrypted candidates in Paneth cells derived from DEG upregulated in WT CH compared to TLR2 cKO CH. X-axis: gene list, Y-axis: absolute Torres score. (**j**) Distribution of proximal (left) and distal (right) Paneth signatures in Paneth cells of chimeric (CH) WT (blue) vs. TLR2 cKO (orange) mice; Significance was determined by Pearson’s *r* test, \*\**p*<0.01, \*\*\*\**p*<0.0001.

**Extended Data Fig. 5.**
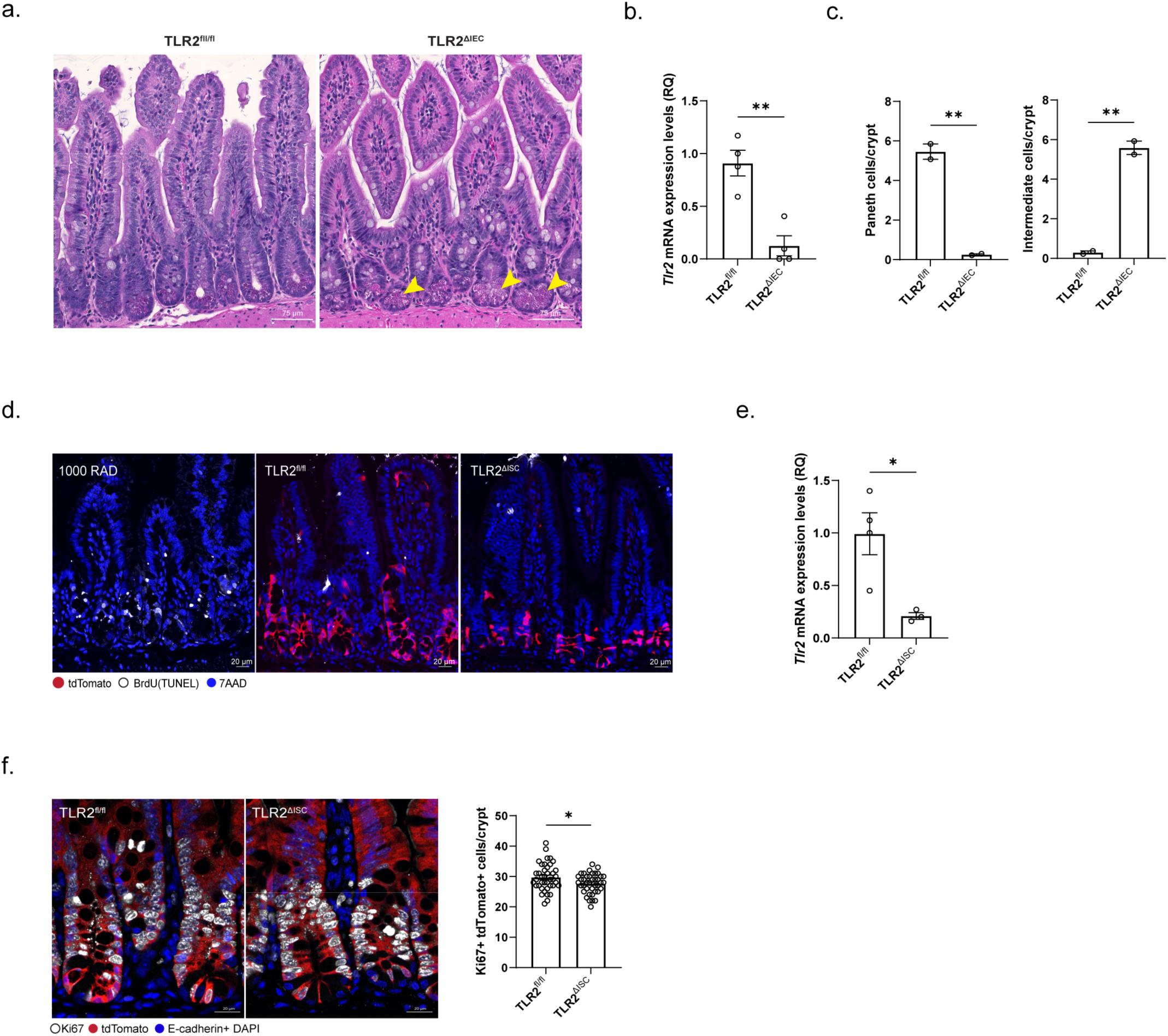
Characterization of epithelial- and stem cell-specific TLR2 deletion. (**a**) Representative H&E images of SI ileum sections from TLR2^fl/fl^ and TLR2^ΔIEC^; scale bars, 75μm; yellow arrows show the changes in granular cells at the bottom of the crypt in TLR2^ΔIEC^ mice. (**b**) Relative quantification (RQ) of *Tlr2* expression in TLR2^fl/fl^ and TLR2^ΔIEC^ mice using qPCR. Data are presented as mean ± SEM. *n*=4 mice per group, \*\**p* <0.01, two-tailed Student’s t-test. (**c**) Goblet-Paneth intermediate cells accumulation is observed in TLR2^ΔIEC^. The number of Paneth (left) and goblet-Paneth intermediate (right) cells per crypt was quantified (right); 25 crypts per mouse, 2 mice per group; data are mean ± SEM; \*\**p* < 0.01, two-tailed *Student’s t-test*. (**d**) TUNEL assay analysis of TLR2^ΔISC^ SI ileal sections shows no signs of cell death after 48h of TLR2 deletion induction. Crypts of positive control (24h post-irradiation (1000 rad), left), TLR2^fl/fl^ (middle), or TLR2^ΔISC^ (right) mice. IFA of tdTomato cells (red), BrdU (TUNEL, white), and DAPI (blue). Scale bar, 20μm. (**e**) Relative quantification (RQ) of *Tlr2* expression levels in TLR2fl/fl (*n*=4 mice) or TLR2^ΔISC^ (*n*=3 mice) using qPCR (right). Data are presented as mean ± SEM; \**p* <0.05, two-tailed *Student’s t-test*. (**f**) IFA of tdTomato (red), Ki67 (white), E-cadherin and DAPI (blue) of distal SI TLR2^fl/fl^ or TLR2^ΔISC^, 5 days after Tamoxifen induction; scale bar, 20μm. (left panel). Quantification of Ki67+ tdTomato+ cells (Right) per crypt. Data are presented as mean ± SEM. 20 crypts per mouse, 2 mice per group. \**p* < 0.05, two-tailed *Student’s t-test*.

**Extended Data Fig. 6.**
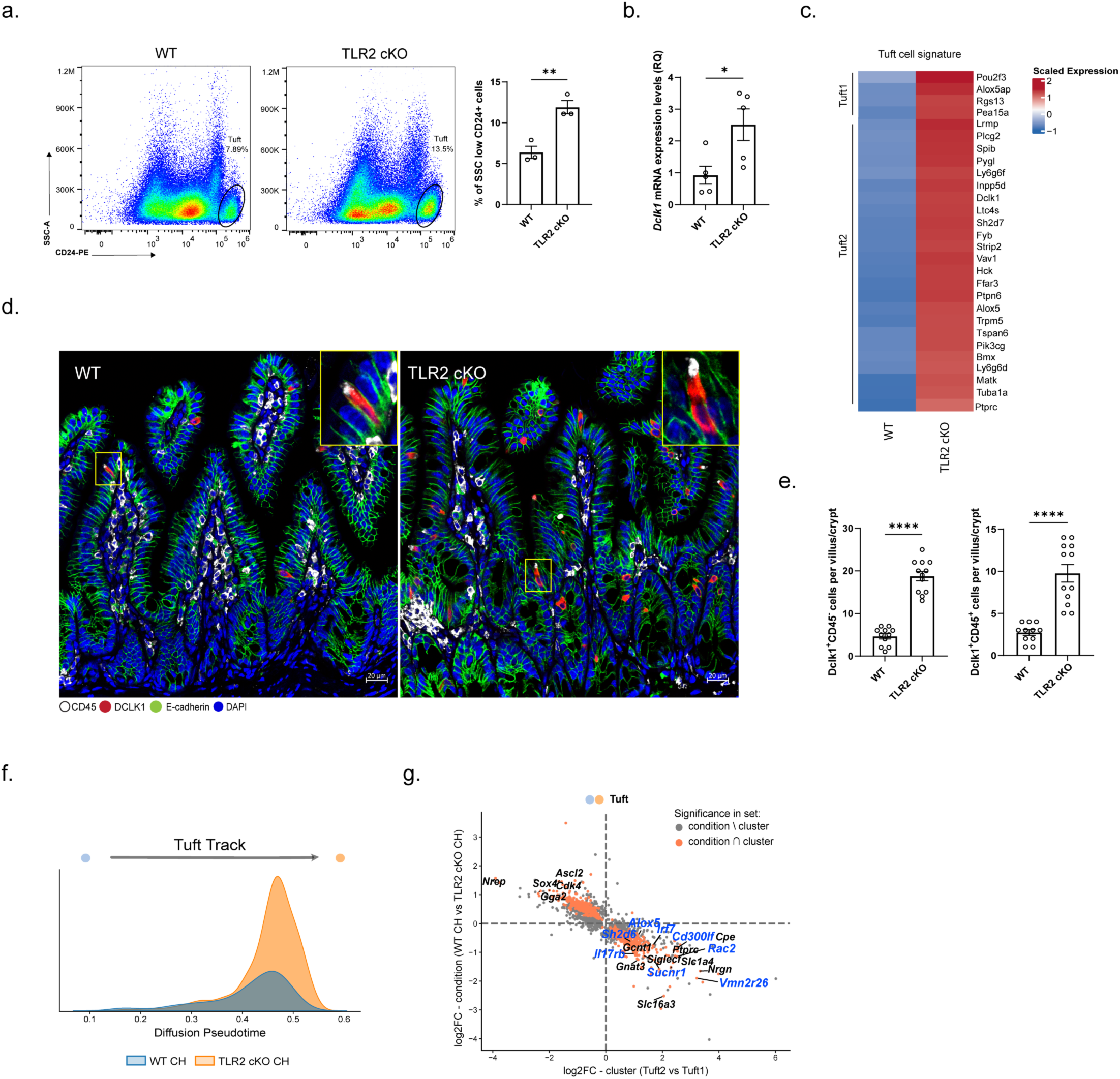
Expansion of microbial-sensing Tuft cells under TLR2 deletion. (**a**) Representative flow cytometry plot (left) and quantification (right) of the percentage of CD24^+^ side-scattered low (SSC^low^) cells among EpCAM^+^ cells in WT vs. TLR2 cKO mice (left panel); quantification of CD24^+^ SSC^low^ percentages in both groups are depicted on a bar graph (right panel); *n*=3 mice per group. Data are presented as mean ± SEM. \*\**p*<0.01, two-tailed *Student’s t-test*. (**b**) Relative quantification (RQ) of *Dclk1* expression levels in EpCAM^+^ cells from WT or TLR2 cKO using qPCR. Data are presented as mean ± SEM. *n*=5 mice per group, \**p* <0.05, two-tailed *Student’s t-test*. (**c**) Heatmap of the top tuft cell markers (Tuft 1 or 2 subsets) in EpCAM^+^ cells from WT (*n*=3) vs. TLR2 cKO (*n*=2) mice out of the DEG using bulk RNA-seq. Average relative expression (mean Z score of log_2_(TPM+1), color bar). (**d-e**) elevation in Tuft 1 and 2 in TLR2 deletion. (d) IFA of DCLK1 (red), CD45 (white), E-cadherin (green), and DAPI. E-Cadherin (green) stains IEC boundaries; scale bar, 20 μm. Inset, x2 magnification. (e) The number of Tuft 1 (DCLK1^+^CD45^-^) and Tuft 2 (DCLK^+^CD45^+^) cells per villus-crypt axis was quantified. Data are presented as mean ± SEM. 10 fields per mouse, \*\*\*\**p* < 0.0001, two-tailed *Student’s t-test*. (**f-g**) Pseudotime analysis for tuft cell differentiation trajectory in chimeric (CH) WT or TLR2 cKO at single cell resolution. (f) Diffusion pseudotime (*dpt*) score comparing the tuft cell differentiation state of chimeric (CH) WT (blue) and TLR2 cKO (orange). Dpt scores showing a left-shift peak in TLR2 cKO cells, indicating accelerated differentiation compared to WT. (g) Scatter plot comparing log2 fold-change between chimeric (CH) WT or TLR2 cKO (y-axis) in tuft cells and log2 fold-change between Tuft1 or Tuft2 (x-axis) clusters. Each point represents one gene. The colors represent different conditions. Highlighted genes (blue) represent Tuft-2–specific bacterial-sensing receptors.

**Extended Data Fig. 7.**
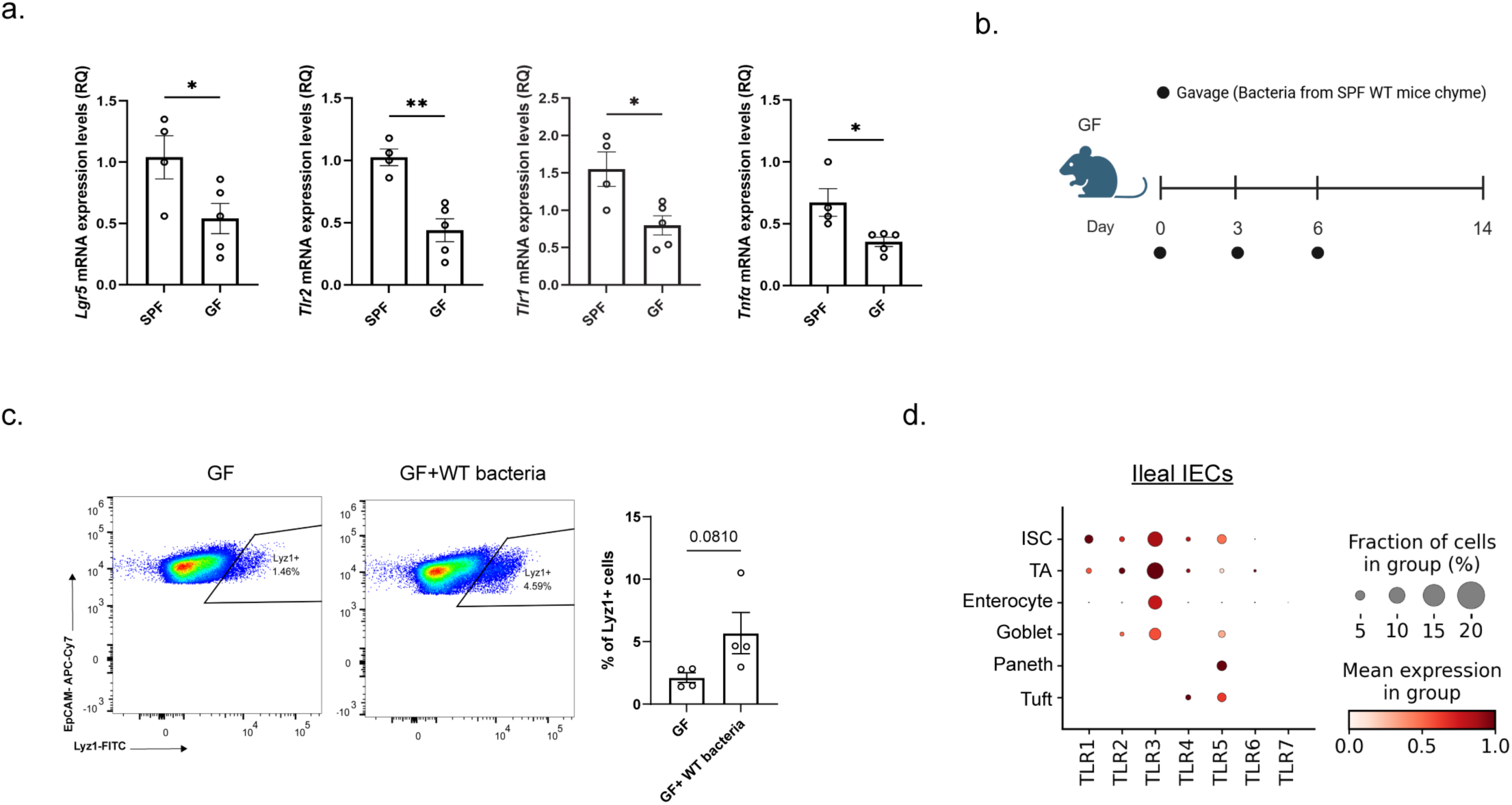
Characterization of the microbiota’s role in TLR2 activation. (**a**) Relative quantification (RQ) of *Lgr5* and TLR2-related genes (*Tlr2, Tlr1 and Tnfα*) expression levels in SPF (*n*=4 mice) or GF (*n*=5 mice) using qPCR (right). Data are presented as mean ± SEM; \**p* <0.05, **p<0.01, two-tailed *Student’s t-test*. (**b-c**) GF colonization experiment. (b) Experimental scheme of colonization experiment in GF mice. (c) Representative flow cytometry plots (left) and quantification (right) of the percentage of Lyz1^+^ cells in ileal crypts of GF (left) or GF colonized with WT microbiota (right). *n*=4 biological repetitions. Data are presented as mean ± SEM; *ns* (non-significant), two-tailed *Student’s t-test*. (**d**) droplet-based epithelial single cell re-analysis of Toll-like receptors (TLRs). Dot plot of 7 human TLRs expressed in different IECs under homeostasis, taken from a publicly available dataset^75^. The dot size indicates the proportion of cells expressing a gene, and the color indicates the mean expression levels. TA, transient amplifying cells.

